# Investigation of fascin1, a marker of mature dendritic cells, reveals a New role for IL-6 signaling in chemotaxis

**DOI:** 10.1101/2020.03.19.979104

**Authors:** Fumio Matsumura, Robin Polz, Sukhwinder Singh, Jürgen Scheller, Shigeko Yamashiro

**Affiliations:** Department of Molecular Biology & Biochemistry, Rutgers University, 604 Allison Rd, Piscataway, NJ 08854; Institute of Biochemistry and Molecular Biology II, Medical Faculty, Universitätsstrasse 1, 40225 Düsseldorf Heinrich-Heine-University Düsseldorf; Department of Pathology & Laboratory Medicine, Medical Science Building (MSB), Rutgers, New Jersey Medical School, 185 South Orange Avenue, Newark, NJ 07101

**Keywords:** IL-6, IL-6Rα, fascin, dendritic cells, chemotaxis, CCR7 internalization

## Abstract

Migration of mature dendritic cells (DCs) to lymph nodes is critical for the initiation of adaptive immunity. While CCR7, a a G-protein-coupled receptor for CCL19/21 chemokines, is known to be essential for chemotaxis of mature DCs, the molecular mechanism linking inflammation to chemotaxis remains unclear. We previously demonstrated that fascin1, an actin-bundling protein, increases chemotaxis of mature DCs. In this paper we showed that fascin1 enhanced Interleukin (IL)-6 secretion and signaling. Furthermore, we demonstrated that IL-6 signaling is required for chemotaxis. Blockage of IL-6 signaling in WT DCs with an anti-IL-6 receptorα (IL-6Rα) antibody inhibited chemotaxis toward CCL19. Likewise, knockout (KO) of IL-6Rα inhibited chemotaxis of BMDCs. The addition of soluble IL-6Rα and IL-6 rescued chemotaxis of IL-6Rα KO BMDCs, underscoring the role of IL-6 signaling in chemotaxis. We found that IL-6 signaling is required for internalization of CCR7, the initial step of CCR7 recycling. CCR7 recycling is known to be essential for CCR7-mediated chemotaxis, explaining why IL-6 signaling is needed for chemotaxis of mature DCs. Our results have identified IL-6 signaling as a new regulatory pathway for CCR7/CCL19-mediated chemotaxis, and suggest that rapid migration of mature DCs to lymph nodes depends on inflammation-associated IL-6 signaling.

## Introduction

Dendritic cells (DCs) are professional antigen presenting cells, playing central roles in adaptive immunity(1, 2). When DCs encounter pathogens, they undergo terminal differentiation, called maturation. Maturation causes massive changes in gene expression as well as morphology and motility, altering the function of DCs from antigen sampling to antigen presentation. While immature DCs move around the peripheral tissues to function as a sentinel, mature DCs respond to chemoattractant stimuli and migrate into lymph nodes as quickly as possible in order to present antigen information to T-cells. Thus, chemotactic migration of mature DCs to lymph nodes is essential for initiating adaptive immunity(3, 4).

Directed migration of mature DCs to lymph nodes requires induction of CCR7, a G-protein coupled receptor, in mature DCs(5–10). CCR7 senses its ligand chemokines (CCL19/21) expressed in lymph nodes(9–11). A gradient of CCL19/21 allows mature DC to migrate from the cell periphery to lymph nodes. The binding of CCL19/21 to CCR7 induces several signal transduction events including activation of MAPKs (ERK1/2 and P38MAPK), PI3K/AKT, tyrosine kinases, Rac, Cdc42 and Rho GTPases(11–16). These downstream signaling events are suggested to be involved in chemotactic migration and survival of mature DCs(12, 13, 17). However, the molecular mechanism linking inflammation to chemotaxis of mature DCs remains unclear.

Fascin1, an actin-bundling protein, is known to promote assembly of filopodia and cell migration(18–20). Fascin1 is induced to a great extent upon DC maturation(21, 22). Fascin1 is absent in immature DCs or other blood cells including B- and T-lymphocytes, macrophages and neutrophils(23). This unique expression pattern suggests that fascin1 plays a unique role in the motility of mature DCs. We found, by analyses of fascin1 KO mice, that fascin1 promotes *in vivo* migration of ear skin DCs into draining lymph nodes after FITC ear painting(24). Fascin1 also increases *in vitro* cell motility and chemotaxis of mature, bone marrow-derived DCs (BMDCs) toward CCL19(24). The importance of fascin1 in cell motility is also observed with other types of cells including normal and transformed cultured cells(25–30).

We previously found that fascin1 KO mature DCs were more susceptible to *Listeria* infection than WT mature DCs(31). Because maturation of DCs triggers secretion of several proinflammatory cytokines including IL-1, IL-6, IL-12 and TNF, the finding has prompted us to examine whether *Listeria* infection alters serum cytokine profiles between WT and fascin1 KO mice. We found that the serum level of IL-6 cytokine is specifically reduced in fascin1 KO mice than in WT mice after *Listeria* infection. We also found that fascin1 KO BMDCs matured by lipopolysaccharide (LPS) showed reduced secretion of inflammatory cytokines including IL-6 and TNFα. Because IL-6 is shown to promote cell motility of a variety of tumor cell lines(32–36), we examined whether fascin1-mediated increase in IL-6 is involved in chemotactic migration of DCs. We found that IL-6 signaling is required for effective chemotaxis of mature DCs toward CCL19, thereby linking IL-6 mediated inflammation to chemotaxis of mature DCs.

## Results

### Fascin1 alters chemokine secretion including IL-6

We analyzed whether infection with *Listeria monocytogenes* alters cytokine secretion profiles in mouse sera. Cytokine profiling was performed using RayBiotech cytokine array analyses (Array C series 2000; see supplemental Fig. 1 for the assignment of 144 unique mouse cytokines of the array 3-5). The analyses revealed a specific reduction in IL-6 levels of fascin1 KO mice in comparison with WT mice (Fig. 1A). Other inflammatory cytokines including IL-10, IL-12, and TGF-β showed no differences between WT and fascin1 KO mice. ELISA assays confirmed that the serum IL-6 level was reduced in fascin1 KO mice (220pg/ml for WT, 50pg/ml for KO) (Fig. 1B). Before infection, no differences in cytokine secretion were observed in sera between WT and fascin1 KO mice (supplemental Fig. 2), suggesting that the change in the serum IL-6 level is associated with immunity.

**Figure 1.**
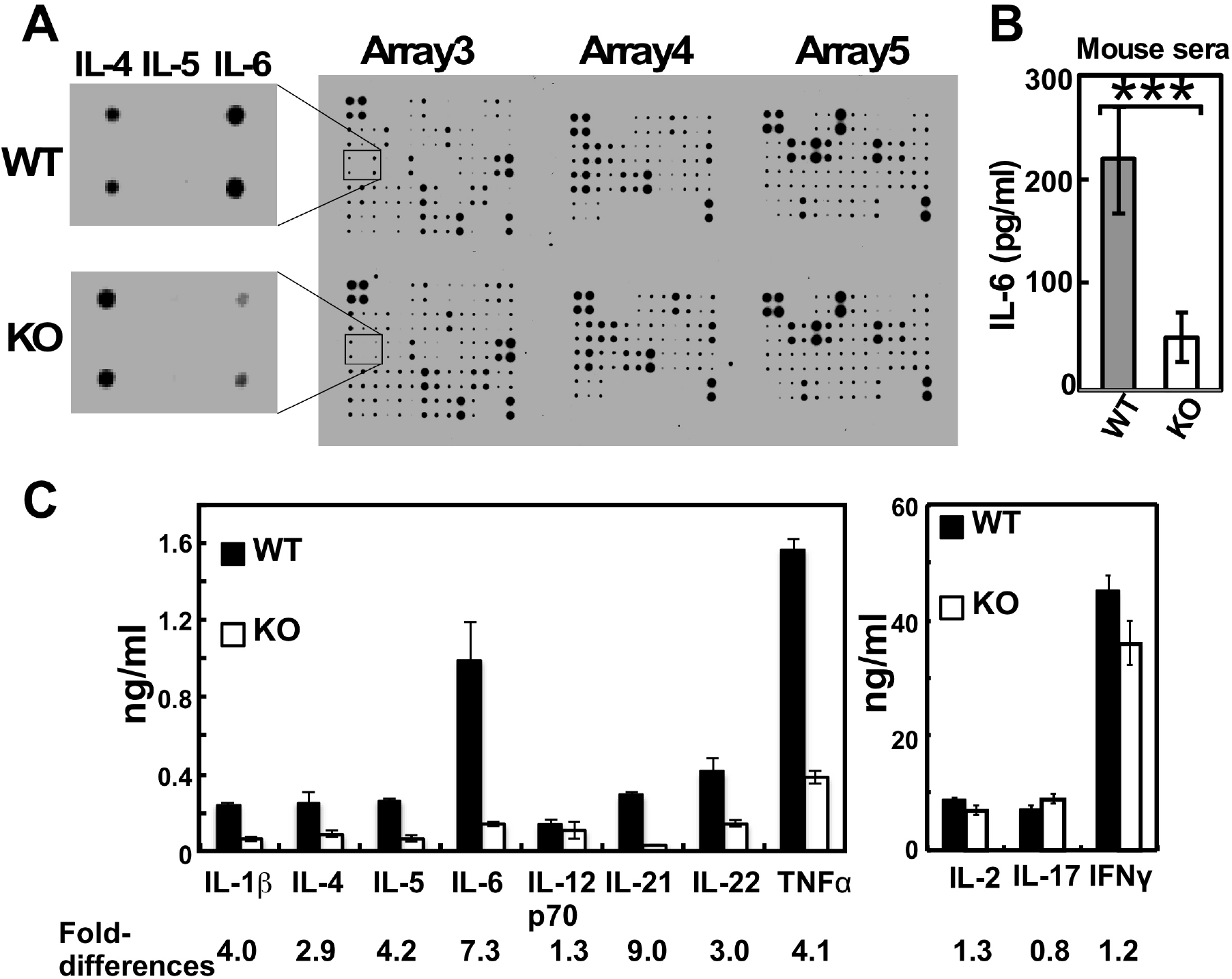
Fascin1 increases IL-6 secretion *in vivo* and *in vitro*. **A**, Analysis of cytokine profiles in sera of WT and fascin1 KO mice 48hr after infection with *Listeria monocytogenes*. Sera were analyzed using RayBio Mouse cytokine antibody Arrays 3-5 (see Supplemental Figure 1 for the positions of each cytokines analyzed). IL-6 is specifically reduced in sera from fascin1 KO mice. **B**, ELISA analyses of IL-6 levels in sera confirmed decreased IL-6 secretion in fascin1 KO mice. ***, p<0.001. Error bars, standard deviation (n=4). **C**, Cytokine profile of culture supernatants of WT and fascin1 KO BMDCs analyzed by Quantibody Mouse Th17 Array1 (RayBiotech, Inc). Fold-differences between WT and KO BMDCs are indicated below each cytokine.

These results indicate that the increase in serum IL-6 levels upon infection depends on immunity-mediated changes in the levels and/or activity of fascin1. DCs induces fascin1 to a great extent upon maturation. Mature DCs are also one of the major sources of IL-6. Therefore, we examined whether fascin1 deficiency altered IL-6 secretion by BMDCs. Quantitative cytokine profiling (RayBio Quantibody mouse Th17 array1) revealed that mature BMDCs from fascin1 KO mice showed reduced secretion of several cytokines including IL1β, IL-4, IL-5, IL-6, IL-21, IL-22, and TNFα (Fig. 1C). No significant changes in the secretion of IL-2, IL-12p70, IL-17 and IFNγ were observed. Among the altered cytokines, IL-6 and IL-21 showed the highest fold-difference between WT and fascin1 KO BMDCs. The reduced IL-6 and TNFα secretion by fascin1 KO BMDCs is consistent with a previous report that fascin1 knockdown by siRNA reduced the secretion of these two cytokines in both human and mouse monocytic leukemia (37) (note that, unlike primary blood cells, these transformed leukemia express fascin1).

### Fascin1 increases IL-6 signaling

The reduced IL-6 secretion is expected to attenuate IL-6 signaling in BMDCs. IL-6 binding to its receptors (IL-6Rα) results in the association between IL-6Rα and gp130, leading to the activation of IL-6 signaling(38). By quantitating the IL-6Rα/gp130 association by Proximity Ligation Assay (PLA), we compared the extents of IL-6 signaling between mature WT and fascin1 KO BMDCs. In PLA assays, the association between two molecules can be detected as a fluorescent speckle.

As Fig. 2A shows, PLA assay revealed that WT mature BMDCs (panel c) exhibit more fluorescent speckles than fascin1 KO mature BMDCs (panel d). The difference is confirmed by the quantitative analyses of fluorescent speckles (Fig. 2B). In contrast to mature BMDCs, immature BMDCs either from WT (Fig. 2A, panel a) or fascin1 KO (Fig. 2A, panel b) showed undetectable levels of PLA signals. The lack of PLA signals in immature BMDCs is consistent with the lack of IL-6 production before DC maturation(39, 40), providing a negative control for the PLA assay. The lower PLA signals in fascin1 KO BMDCs are not caused by decrease in the expression levels of IL-6Rα or gp130 (Supplemental Fig. 3): Both immunofluorescent localization and the expression levels of surface IL-6Rα and gp130 were similar between WT and fascin1 KO BMDCs. Taken together, these results indicate that fascin1 increases IL-6 secretion, leading to higher IL-6 signaling in mature BMDCs.

**Figure 2.**
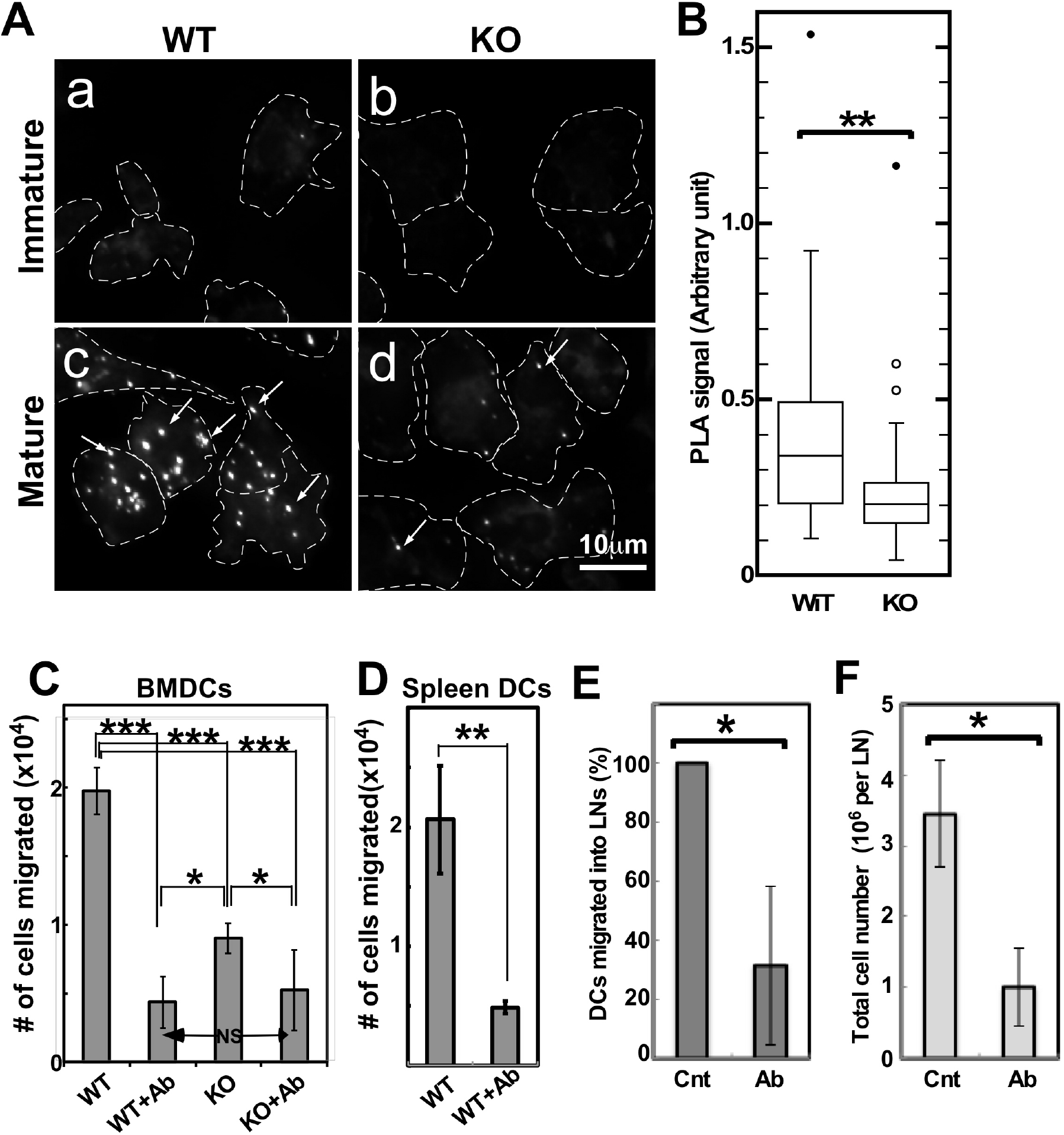
IL-6 signaling is critical for chemotaxis of DCs. **A**, Higher IL-6 signaling of WT BMDCs than that of fascin1 KO BMDCs. Proximity Ligation Assay (PLA) was used to determine the *in situ* association between IL-6Rα and gp130, representing the extent of IL-6 signaling. a & b, immature BMDCs; c & d, mature BMDCs. a & c, WT. b & d, fascin1 KO. Fluorescence speckles (arrows) indicate the association between IL-6Rα and gp130. Cell boundaries are indicated by dashed lines. **B**, Quantitative analyses of PLA signals in WT and fascin1 KO BMDCs (n=51). **, p<0.01. **C**, Effects of the inhibition of IL-6 signaling on chemotaxis of BMDCs. Collagen-coated, modified Boyden chamber chemotaxis assays were used to determine chemotaxis of WT BMDCs (WT, n=6); WT with a neutralizing antibody against IL-6Rα (W+Ab, n=3); fascin1 KO BMDCs (KO, n=6); KO with a neutralizing antibody (KO+Ab, n=3). Error bars are standard deviations. **D**, A neutralizing antibody against IL-6Rα also inhibits chemotaxis of mature DCs isolated directly from spleens (n=3). Collagen-coated modified Boyden chamber chemotaxis assay was used. **E** & **F**, Effects of a neutralizing antibody on *in vivo* chemotaxis (E) and cellularity (F). CSFE-labeled BMDCs together with a control or neutralizing antibody were injected into the dorsal part of the foot. After 24hr, BMDCs migrated into draining lymph nodes were counted by flow cytometry (E). Total cell numbers of lymph nodes were also counted to show differences in cellularity (F).

### IL-6 signaling is critical for chemotaxis of mature DCs

IL-6 signaling has been reported to increase cell motility of a variety of tumor cell lines(32–36). We thus hypothesized that fascin1-mediated, higher IL-6 signaling may account for higher chemotaxis of WT BMDCs in comparison with fascin1 KO BMDCs. To test this hypothesis, we added a neutralizing IL-6Rα-antibody to WT and fascin1 KO BMDCs just before activation with LPS, and examined, by modified Boyden chamber assays, whether the blockage of IL-6 signaling affects chemotaxis.

We found that IL-6 signaling is required for effective chemotaxis of mature BMDCs (Fig. 2C). The addition of a neutralizing IL-6Rα-antibody greatly reduced chemotaxis of WT BMDCs to one-fifth of that of control. Consistently with the previous result(24), fascin1 KO reduced chemotaxis to one-half of the level shown by WT BMDCs. The addition of an IL-6Rα antibody to fascin1 KO BMDCs further reduced chemotaxis to a very low level similar to that of IL-6Rα-inhibited WT BMDCs. The inhibition of chemotaxis is not due to blockage of DC maturation: The addition of the neutralizing antibody did not significantly alter the expression of maturation markers of WT BMDCs (Supplemental Fig. 4A-E): We further found that the IL-6Rα antibody was able to block chemotaxis even when the antibody was added 16hr after LPS addition, indicating that sustained IL-6 signaling is required for chemotaxis of mature BMDCs. These results suggest that chemotaxis depends on the strengths of IL-6 signaling.

A recent study using gene expression analyses suggests that BMDCs generated in the presence of GM-CSF may not represent any subtype of *in vivo* DCs(41). While this issue is still under debate(42–47), it is important to determine whether the effect of IL-6 signaling on chemotaxis is observed with DCs directly isolated from tissues. Therefore, we examined whether chemotaxis of spleen DCs from WT mice depends on IL-6 signaling. As Fig. 2D shows, the blockage of IL-6 signaling inhibited chemotaxis of spleen DCs. This result suggests that IL-6 signaling is generally required for chemotaxis of conventional mature DCs.

We further examined whether inhibition of IL-6 signaling affects DC chemotaxis *in vivo* in mice. Mature BMDCs were labeled with CSFE and mixed with a neutralizing IL-6Rα antibody or with a control antibody. They were then injected into a dorsal part of foot. Twenty-four hours later, CSFE-labeled BMDCs migrated into draining lymph nodes were counted by flow cytometry. As Fig. 2E shows, the number of BMDCs migrated into lymph nodes was greatly reduced when a neutralizing IL-6Rα antibody was co-injected. The total cell number of lymph nodes was also reduced in the presence of the neutralizing IL-6Rα antibody (Fig. 2F), confirming that DC did not reach to the lymph nodes for activation. These results suggest that IL-6 signaling is critical for *in vivo* migration of mature DCs toward draining lymph nodes.

### Fascin1/IL-6 signaling axis controls the directionality of DC chemotaxis

Our results indicate that fascin1 promotes IL-6 signaling, which is critical for chemotaxis of mature DCs. Chemotaxis depends on speeds and/or directness of migration. To determine which factor(s) fascin1 and IL-6 signaling affect, we performed live cell imaging of BMDCs migrating toward a CCL19 gradient in a collagen gel as described(48). We found that the lack of fascin1 and the inhibition of IL-6 signaling both reduce directionality of DC migration without altering migration speeds.

Figs 3A-a and c show the migration tracks of WT and fascin1 KO BMDCs, respectively. These tracks showed that WT BMDCs (Fig. 3A, panel a, see supplemental videos 1a and b) apparently moved more consistently along a CCL19 gradient (shown by arrows) than did fascin1 KO BMDCs (Fig 3A, panel c, supplemental videos 2a and b). Rose plot diagrams (Fig. 3A panels b and d; an angular histogram of the fractions of cells moving toward each direction with a bin of 10 degree) supports the above notion: More WT cells moved along the direction of the chemokine gradient than KO cells. The blockage of IL-6 signaling also reduced directed migration (Fig. 3A-e, supplemental videos 3a and b): The migration tracks of WT BMDCs in the presence of the neutralizing antibody against IL-6Rα migrated randomly, which is obvious with the Rose plot analysis (Fig. 3A-f). These analyses suggest that both the deficiency of fascin1 and the blockage of IL-6 signaling reduced directional migration.

**Figure 3.**
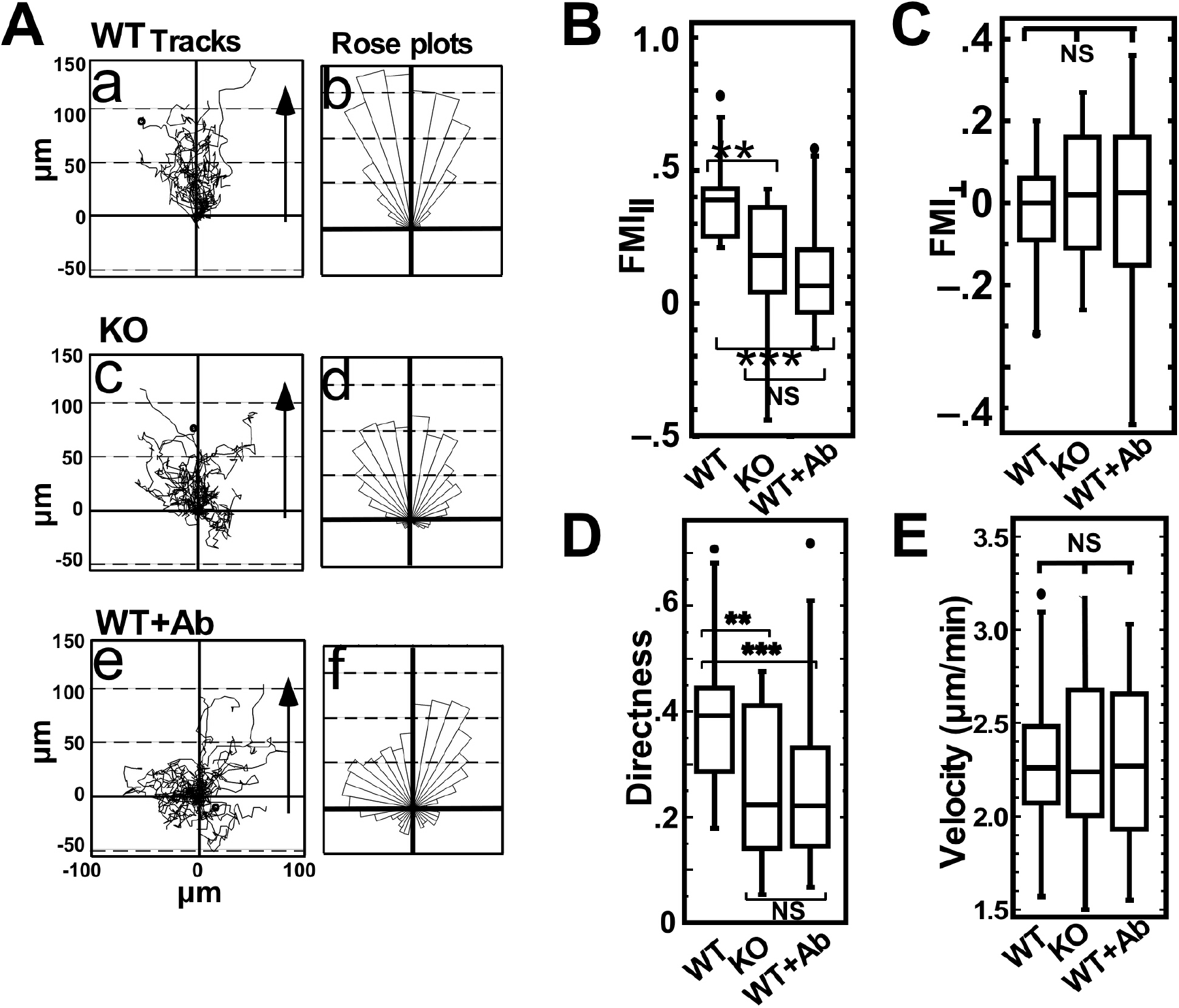
IL-6 signaling is required for directional migration of BMDCs. A, Analyses of migration tracks. Live cell imaging of BMDCs migrating in a collagen gel toward CCL19 was performed to obtain migration tracks (a, c & e) and rose diagram plots for directionality(b, d & f). a & b, WT (n=19): c& d, fascin1 KO (n=23): e & f, WT BMDCs in the presence of a neutralizing antibody against IL-6Rα (n=28). WT BMDCs (a & b) showed more consistent migration toward CCL19 than fascin1 KO counterparts (c & d). Blockage of IL-6 signaling inhibits directed migration of WT BMDCs (e & f). Arrows, the direction of a CCL19 gradient. B-E, Box plot analyses of parallel (B, FMI∥) and perpendicular (C, FMI⊥) forward migration indexes, directness (D), and migration speeds (E). *, p<0.05; **, p<0.01; ***, p<0.001; NS, not significant. The experiments were performed at least 3 times for each condition and a representative result is given.

To obtain more quantitative data of the directionality (i.e. efficiency of chemotactic migration), we defined two parameters of FMI_∥_ and FMI_⊥_ as described(49, 50), which represent the forward migration of cells parallel and perpendicular to the chemoattractant gradient, respectively (see supplemental Fig. 5). We chose the coordinates so that the x-axis is perpendicular to the direction of the gradient and the y-axis is parallel to the gradient. The FMI_⊥_ and FMI_∥_ value were derived after dividing the (x, y) coordinates of each migrating cell at the endpoint by the migration path length, respectively(49, 50). Positive FMI_∥_ values indicate migration toward the chemoattractant source, while negative values indicate migration away from the chemoattractant source. The higher FMI_∥_ values show the more efficient migration toward the chemoattractant source.

As Fig. 3B, panel a, shows, WT BMDCs show a higher median value (0.39) of FMI_∥_ (forward migration index parallel to the gradient) than that (0.18) of KO cells (p=0.0023). The addition of an anti-IL-6Rα neutralizing antibody to WT BMDCs also reduced the median value of FMI_∥_ from 0.39 (control) to 0.07 (p less than 0.0001). The median values of the perpendicular migration index (FMI_⊥_), on the other hand, are all near zero for WT, fascin1 KO and IL-6Rα-blocked WT BMDCs (Fig. 3C), which is consistent with the absence of a chemotactic gradient in the perpendicular direction.

We also measured directness of migration (Fig. 3D), which is often analyzed for chemotactic migration. Directness (sometimes confusingly called directionality) is a scalar quantity that measures how straight the cells travel in a given distance though it does not necessarily indicate movements toward the chemoattractant (supplemental Fig. 5). The median values of directness are 0.39 for WT; 0.22 for KO (p=0.0036, compared with WT) and 0.19 for IL-6Rα-inhibited WT (p=0.0005, compared with WT). The difference between fascin1 KO and IL-6Rα-inhibited WT is not statistically significant.

Directed migration also depends on migration speeds. We quantified the average migration speeds of each cell moving in a collagen gel (Fig. 3E). The speeds of WT, fascin1 KO BMDCs and IL-6Rα-blocked WT BMDCs were found to be statistically similar. This observation differs from our previous result that WT BMDCs moved faster than fascin1 KO migration on 2D surface(24). The discrepancy is most likely due to the difference in the mode of migration between 2D and 3D. On a 2D surface, WT, mature BMDCs were able to disassemble adhesive structure called podosomes, resulting in higher migration speeds(24). In a 3D gel, DCs are known to show adhesion-independent amoeboid movements(51).

Taken together, these results indicate that IL-6 signaling is critical for the directionality of chemotactic migration toward CCL19 without altering migration speeds, and suggest that the reduced IL-6 secretion by fascin1 KO is likely to explain the inhibition of chemotaxis.

### BMDCs from IL-6Rα KO mice show impaired chemotaxis toward CCL19

The above results predict that knockout of the IL-6Rα gene would reduce chemotaxis of mature DCs toward CCL19. To address this prediction, we compared chemotaxis of BMDCs from IL-6Rα KO mice and their heterozygous littermate. The lack of IL-6Rα expression in IL-6Rα KO BMDCs was confirmed by immunofluorescence (Fig 4A).

**Figure 4.**
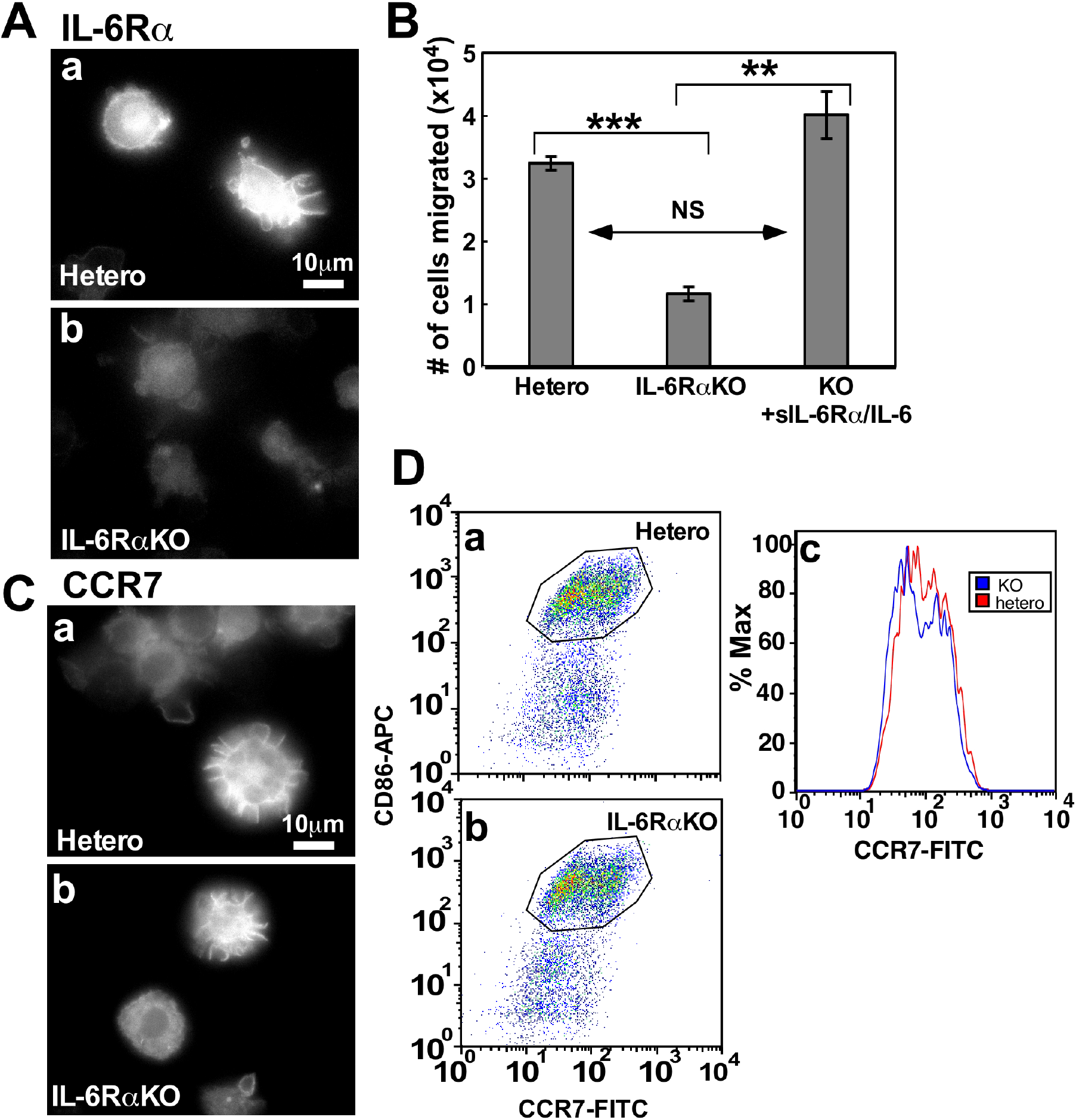
Impaired chemotaxis of IL-6Rα KO BMDCs and its restoration by the addition of soluble IL-6Rα (sIL-6Rα) and IL-6. BMDCs were isolated from heterozygous (hetero) and IL-6Rα KO mice. **A**, Immunofluorescence of IL-6Rα in heterozygous (a) and IL-6Rα KO BMDCs (b) confirming the lack of IL-6Rα in KO BMDCs. **B**, Collagen-coated, modified Boyden chamber chemotaxis assays of heterozygous (hetero), IL-6Rα KO, and IL-6Rα KO in the presence of sIL-6Rα and IL-6 (KO+sIL-6R/IL-6). Note that KO BMDCs show reduced chemotaxis, which was rescued by the addition of soluble IL-6Rα (sIL-6Rα) and IL-6. **C** & **D**, IL-6Rα KO did not alter surface expression of CCR7. Immunofluorescence (C) showing similar levels of surface CCR7 in heterozygous (a) and IL-6Rα KO BMDCs (b). Flow cytometry analyses (D) confirming that IL-6Rα KO did not affect the levels of CCR7 surface expression. After CD86-positive heterozygous (a) and IL-6Rα KO (b) BMDCs were gated, the levels of CCR7 surface expression were examined in histogram (c).

Modified Boyden chamber chemotaxis assays revealed that BMDCs from IL-6Rα KO mice show much lower chemotaxis than BMDCs from heterozygous mice (Fig. 4B). To examine whether the lack of IL-6 signaling is indeed responsible for reduced chemotaxis, we performed rescue experiments by the addition of both soluble IL-6Rα (sIL-6Rα) and IL-6. IL-6 binds to sIL-6Rα, the complex of which is associated with membrane-bound gp130, inducing IL-6 signaling (trans-signaling)(52–56). We found that the addition of IL-6 and sIL-6α completely rescued the impaired chemotaxis of IL-6Rα KO BMDCs, underscoring the critical role of IL-6 signaling in chemotaxis of mature DCs.

The reduced chemotaxis of IL-6Rα KO BMDCs could be due to a change in CCR7 expression. To exclude this possibility, we compared surface expression of CCR7 between KO and heterozygous BMDCs. As Fig. 4C shows, immunofluorescence revealed that IL-6Rα KO did not affect CCR7 surface expression. Flow cytometry analyses (Figs 4D & E) confirmed that WT and IL-6Rα KO BMDCs express similar levels of CCR7.

### IL-6 signaling controls ligand-induced internalization of CCR7

Recycling of chemokine receptors including CCR7 has been shown to control chemotaxis(57, 58). We examined whether inhibition of IL-6 signaling affects recycling of CCR7. We found that IL-6 signaling is essential for receptor internalization, the first step for receptor recycling. In control BMDCs (Fig. 5A, panel a), CCR7 showed rapid internalization when CCL19 was added. In the presence of a neutralizing IL-6Rα antibody, however, CCR7 internalization is almost completely blocked (panel b). These results suggest that IL-6Rα KO BMDCs should also show impaired CCR7 internalization. As Fig. 5B, panel a, shows, CCL19-induced, CCR7 internalization was also blocked with IL-6Rα KO BMDCs. Importantly, the addition of soluble IL-6Rα and IL-6 restored CCR7 internalization (panel b), reinforcing the notion that IL-6 signaling is critical for CCL19-induced internalization of CCR7.

**Figure 5.**
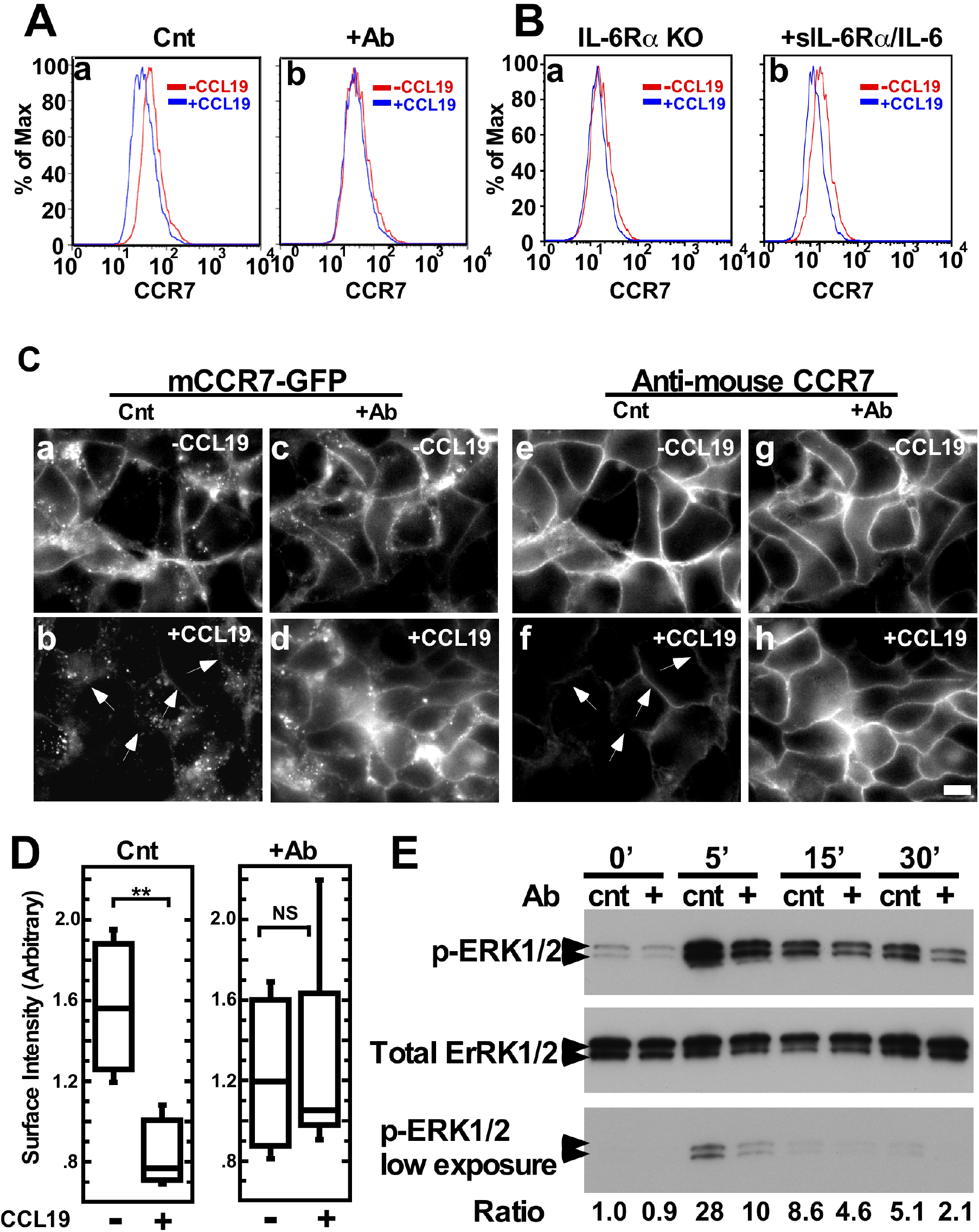
IL-6Rα signaling is critical for CCR7 internalization and CCL19-induced ERK1/2 phosphorylation. **A**, Effects of the blockage of IL-6 signaling on CCR7 internalization. WT BMDCs were stimulated with CCL19 for 5 min in the presence of either control (a) or neutralizing antibody (b), and then surface CCR7 expression was determined by flow cytometry. Surface expression of CCR7 was examined with CD86^+^ BMDCs before (-CCL19) and after CCL19 addition (+CCL19). Note that CCR7 internalization was blocked in the absence of a neutralizing antibody to IL-6Rα. **B**, Effects of IL-6 signaling on CCR7 internalization of IL-6Rα KO BMDCs. IL-6Rα KO BMDCs were stimulated with CCL19 in the absence (a) or presence of sIL-6Rα (soluble IL-6Rα) and IL-6 (b). Note that IL-6Rα KO BMDCs showed impaired CCR7 internalization (a), which was rescued by the addition of sIL-6Rα and IL-6 (b). **C**, Inhibition of CCR7 internalization by the blockage of IL-6 signaling in HEK293T epithelial cells. HEK293T cells stably expressing mouse CCR7-GFP were stimulated with CCL19 in the presence of either control or neutralizing antibody against IL-6Rα. a-d, fluorescence images of CCR7-GFP in control cells (a & b) or IL-6Rα neutralizing antibody-treated cells (c & d) before (a & c) or after addition of CCL19 (b & d). The same cells were counter stained with an antibody specific to mouse CCR7 to detect only surface CCR7 (e-h) in control (e & f) or IL-6Rα neutralizing antibody-treated cells (g & h) before (e & g) or after (f & h) addition of CCL19. **D**, Quantitative measurements of surface CCR7 immunofluorescence (panels e-h of C) at cell-to-cell contacts. In contrast to the control, the addition of a neutralizing antibody against IL-6Rα blocked internalization of CCR7. **E**, Effects of the blockage of IL-6 signaling on CCL19-induced ERK1/2 phosphorylation of WT BMDCs. WT BMDCs were stimulated with CCL19 in the presence of a control or neutralizing antibody, then ERK1/2 phosphorylation levels were determined at 0, 5, 15 and 30min after CCL19 addition using Western blotting with a phosphospecific ERK1/2 antibody. For normalization, the same membranes were reblotted with a pan ERK1/2 antibody. The ratios of phosphoERK1/2 to total ERK1/2 are indicated below the figure.

The inhibition of CCR7 internalization by the blockage of IL-6 signaling was also observed with HEK293T cells that stably expressed a mouse CCR7-GFP (Fig. 5C). Before CCL19 addition, CCR7-GFP was mostly localized at cell-to-cell contacts (panel a). When CCL19 was added, CCR7-GFP at cell-to-cell contacts was greatly reduced with concomitant increase of GFP signals in the cytoplasm (panel b), indicating internalization of CCR7-GFP. In the presence of a neutralizing IL-6Rα antibody, however, CCR7-GFP signals remained at cell-to-cell contacts with much less cytoplasmic signals, indicating the blockage of the internalization of CCR7 (compare panel c with d, Fig. 5C).

The blockage of internalization could be clearly observed when surface CCR7 was visualized. To do so, the HEC293T cells were stained with mouse-specific antibody against CCR7 (clone 4B12) without permeabilization. As expected, this antibody detected only surface CCR7 at cell-to-cell contacts. Without an anti-IL-6Rα neutralizing antibody, CCR7 staining at the cell-to-cell contacts is largely lost by the addition of CCL19 (compare panel e with f). In the presence of the neutralizing antibody, on the other hand, the staining at the cell-to-cell contacts remained largely unchanged (compare panel g with h). Quantitative measurements of fluorescence intensities at the cell-to-cell contacts supported this notion (Fig. 5D).

Internalization of CCR7 occurs via clathrin-coated endocytosis, which are known to be coupled with the activation of ERK1/2 signaling(59). We thus examined whether an IL-6Rα neutralizing antibody affects ERK1/2 phosphorylation. Fig. 5E shows the time courses of ERK1/2 phosphorylation in BMDCs in the presence of a control or a neutralizing IL-6Rα antibody before and 5, 15 and 30min after the addition of CCL19. The ratios of phospho-ERK1/2 to total ERK1/2 at each time point were presented at the bottom of the figure. The neutralizing antibody reduced ERK1/2 phosphorylation throughout the entire time course: the level of ERK1/2 phosphorylation was reduced to one-third at 5min, and to about half at both 15 and 30min time points. Taken together, these results suggest that IL-6 signaling is required for CCR7 internalization, resulting in lowering ERK1/2 phosphorylation, one of the CCL19-mediated CCR7 downstream events.

The inhibition of CCR7 internalization by the blockage of IL-6 signaling could be due to lower binding between CCR7 and CCL19. To examine whether inhibition of IL-6 signaling interferes with binding of CCR7 to CCL19, we used a CCL19-human IgG Fc fusion protein as a ligand(60, 61) and the binding was detected by labeling with fluorescently-labeled anti-human IgG. The levels of the CCL19 association were not altered with or without the addition of the neutralizing IL-6Rα antibody (supplemental Fig. 4F). These results indicate that inhibition of IL-6 signaling did not alter binding of CCL19 to CCR7.

### Fascin1 increases the level of mRNA of IL-6

Finally, we examined whether fascin1 affects IL-6 mRNA levels. We used PrimeFlow RNA assays (Affymetrix), in which transcript levels are quantitated by single molecule fluorescent *in situ* hybridization (FISH) following amplification. As Fig. 6A shows, IL-6 mRNA levels were higher in wild type BMDCs (panel c) than in fascin1 KO (panel d) BMDCs when they were matured with LPS overnight. Immature BMDCs from WT (panel a) or fascin1 KO (panel b) showed very low levels of IL-6 mRNA, which is consistent with the lack of IL-6 secretion by DCs before maturation(39, 40). Quantitative measurements of *in situ* mRNA signals confirmed that WT BMDCs showed higher mRNA levels than fascin1 KO counterparts (Fig. 6B). This result is not consistent with the previous report that fascin1 increased translational efficiency by favoring polysome formation without altering IL-6 mRNA levels(37): While the exact reason for this discrepancy is unknown, our finding suggests that fascin1 may enhance transcription of the IL-6 gene. Alternatively, fascin1 may stabilize IL-6 mRNA as cytokine mRNAs including IL-6 and TNFα mRNAs show regulated stability upon cellular activation(62–66).

**Figure 6.**
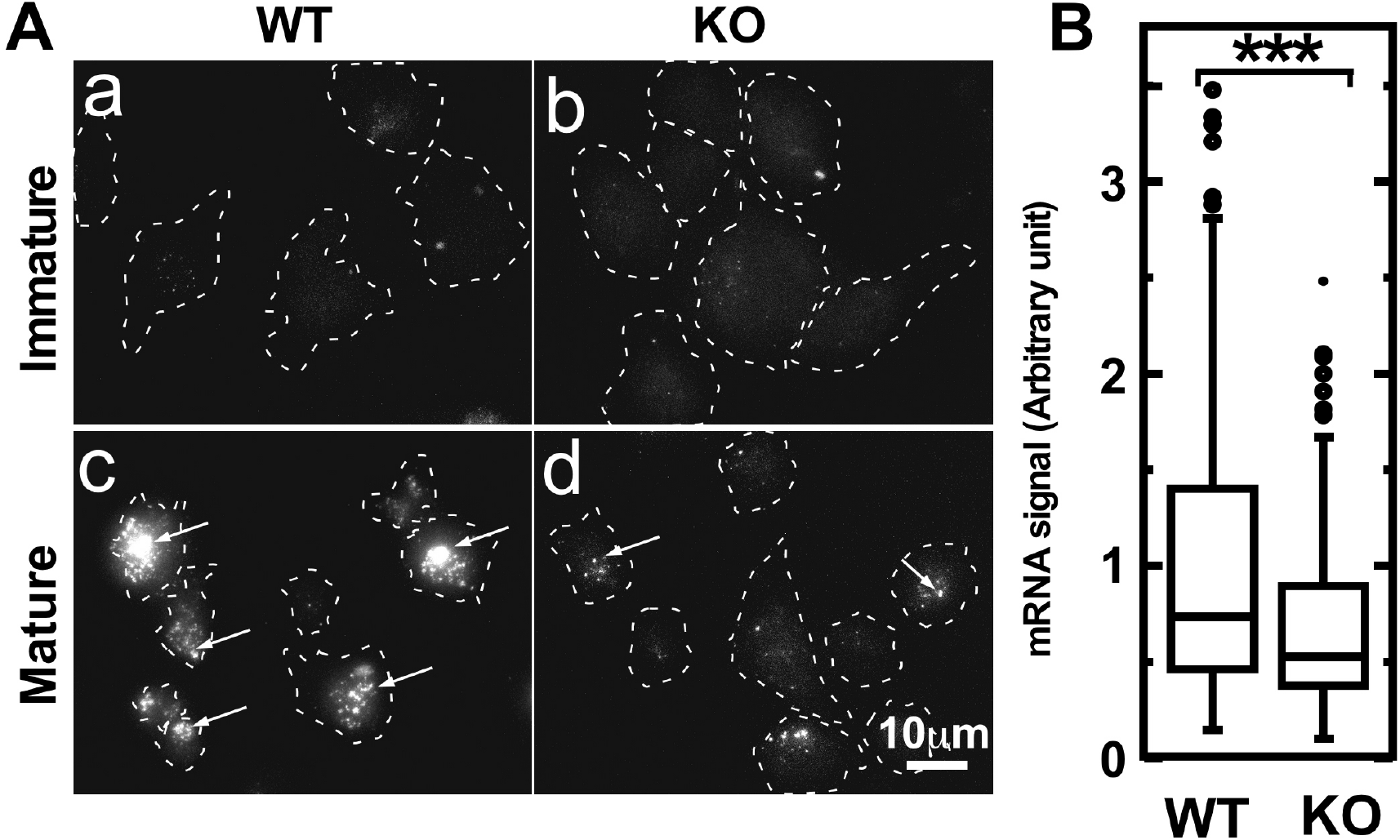
WT BMDCs show higher levels of IL-6 mRNA. A, PrimeFlow RNA assay to determine *in situ* IL-6 mRNA levels in WT (a & c) and fascin1 KO (b & d) BMDCs. a & b, immature BMDCs; c & d, mature BMDCs. Fluorescence dots (indicated by arrows) represent *in situ* hybridized and amplified IL-6 mRNA. Mature wild type BMDCs (c) show more and higher levels of fluorescent dots than mature fascin1 KO BMDCs (d). Note that immature BMDCs either from WT (a) or KO (b) show very low mRNA levels, which is consistent with the lack of IL-6 secretion in immature BMDCs. Dashed lines, Cell boundaries. B, Quantitative analysis of in situ mRNA signals of mature wild and KO BMDCs (>200 cells).

## Discussion

### Critical roles of IL-6 signaling in CCR7 internalization and chemotaxis of mature DCs

We have found that IL-6 signaling is necessary for CCL19-induced internalization of CCR7. This finding could explain why IL-6 signaling is required for directional migration of DCs along a CCL19 chemoattractant gradient. The binding of CCL19 to CCR7 initiates ligand-induced internalization of CCR7 via clathrin-dependent recycling endocytosis(67). The internalized CCR7-CCL19 complexes are then dissociated in the endosomes where these two molecules take two endocytic pathways(67): Whereas CCL19 is degraded in lysosomes, CCR7 is recycled back to the plasma membrane, allowing recycled CCR7 ready for “second round” signaling with a new CCL19 ligand. Therefore, the blockage of CCR7 internalization would greatly decrease CCR7 signaling for sensing CCL19 gradient, resulting in lower chemotaxis.

This notion is consistent with a previous report(58). A ubiquitination-deficient mutant of CCR7 showed impaired internalization in a steady state, reducing an intracellular pool of CCR7. This led to impaired recycling upon CCL19 binding, resulting in the inhibition of chemotaxis toward CCL19(58). A similar role of receptor internalization on chemotaxis was also observed with CXCR2(57): A CXCR2 mutant with a mutation in its LLKIL motif at the C-terminus is unable to undergo ligand-induced internalization and show impaired chemotaxis(57). It should be noted, however, that ligand-induced receptor internalization is not required for all types of GPCR-mediated chemotaxis: An internalization-deficient mutant of CCR2B has been shown to support chemotaxis toward a MCP-1 gradient(68).

What could be a mechanism by which IL-6 signaling controls CCR7 internalization? Internalization of CCR7 starts when CCR7 is associated with β-arrestin(69, 70). This association depends on the phosphorylation of the cytoplasmic C-terminus of CCR7 by a kinase like GRKs (G protein-coupled receptor kinases) (70, 71). CCR7-β-arrestin complexes bind to adaptin, leading to the assembly of clathrin-coated pits for endocytosis of CCR7. IL-6 signaling may affect any of these events. Depending on which step IL-6 signaling controls, the blockage of IL-6 signaling could have distinct effects on CCR7 desensitization. For example, if IL-6 signaling is required for the CCR7-β-arrestin association, then the blockage of IL-6 signaling would inhibit desensitization of CCR7. On the other hand, if IL-6 signaling control the β-arrestin-adaptin association, then the blockage of IL-6 signaling would keep CCR7 desensitized. We are in the process of determining these possibilities.

### Inflammation-dependent and independent migration of CCR7-mediated chemotaxis

We have demonstrated that chemotaxis of mature DCs toward CCL19 depends on IL-6 signaling, which is associated with inflammation. However, this dependence on IL-6 signaling does not apply for all types of CCR7-mediated chemotaxis. For example, naïve T-cells express CCR7, which is critical for their homing to lymph nodes. Their migration occurs without inflammation, and therefore should not require IL-6 signaling. This IL-6-independent migration of T-cells is consistent with the lack of fascin1 in T-cells. Thus, the mechanism of T-cell homing to lymph nodes must be very different from chemotaxis of mature DCs.

The inflammation-independent chemotactic migration may also explain why more DCs were found in lymph nodes of IL-6 KO mice than in those of WT counterparts(72). The increase of DCs in IL-6 KO mice was observed at a steady-state condition without inflammation. Migrating DCs in the steady state may, at least in part, represent DCs “semi-matured” by a self-antigen: It has been reported that these “semi-matured” DCs express CCR7 and migrate into lymph nodes, establishing peripheral tolerance(11). Such migration may not require rapid migration as observed with “fully-matured” DCs during pathogen-induced inflammation. “Fully matured” DCs, on the other hand, need to migrate into lymph nodes as quickly as possible for the initiation of adaptive immune responses against invading pathogens. We suggest that IL-6 signaling is required to ensure that migration of “fully-matured” DCs is inflammation-dependent.

### Possible role of IL-6 signaling in lymphatic metastasis of cancer cells

A number of studies showed that fascin1 promotes metastasis of many cancers including breast, colorectal and gastric carcinoma(28, 29, 73–78). Fascin1 siRNA or fascin1 inhibitors have been reported to inhibit cancer cell migration *in vitro*, as well as metastasis in an *in vivo* mouse model(26–30). These results have been interpreted in the following way: Fascin1 promotes filopodia assembly, thereby accelerating cancer cell migration(18, 24, 25, 79).

Our current results suggest a new role of fascin1 in cancer cell migration via enhancing IL-6 signaling. IL-6 signaling has been reported to increase migration of several types of cancer cells(36, 80–86). Thus, upregulation of fascin1 observed in many cancer cells is likely to simultaneously enhance IL-6 signaling, thereby promoting cancer cell migration. CCR7 expression is often observed for cancer cells with lymphatic metastasis(87). It is possible that fascin1 upregulation in such cancer cells promotes migration to nascent lymph nodes by enhancing IL-6 signaling. The fascin1/IL-6 signaling axis could be a target of therapeutic treatments for lymphatic metastasis of CCR7-expressing cancer cells.

## Methods and Materials

### Reagents, antibodies and mice

The primary antibodies used were: APC- (cat #,17-0862-81) and FITC-conjugated rat anti-CD86 (11-0862-81), AF647-labeled rat anti-mouse IL-6 (11-7061-81) and unconjugated rat anti-mouse CCR7 antibody (16-1971-85, clone 4B12, all from eBioscience, San Diego, CA); FITC-labeled MHC-II (107605), FITC-labeled CD80 (104705), and unconjugated anti-mouse IL-6Rα (115807, all from Biolegend, San Diego, CA); goat anti-mouse IL-6Rα (AF1830, R&D systems, Minneapolis, MN); rabbit anti-gp130 (sc-9045, Santa Cruz Biotech, Dallas, TX); and rabbit anti-CCR7 polyclonal antibody (TA310252, Origene, Rockville, MD). The secondary antibodies were FITC-, Cy3-, or AF647-labeled anti-rat Fab fragment and FITC- or Cy3-labeled anti-goat or anti-rabbit antibodies (Jackson ImmunoResearch, West Grove, PA). GM-CSF and CCL19 (MIP3β) were purchased from Invitrogen (Carlsbad, CA) and Miltenyi Biotec (Auburn, CA).

Fascin1 KO mice were prepared as previously reported(88). For each experiment with homozygous mice, their WT littermates were used as a control whenever possible. IL-6Rα KO mice(89) were obtained from Jackson Laboratory and maintained at Heinrich-Heine University of Düsseldorf. WT or heterozygous littermate mice were used as a control. No phenotypic differences were detected between WT and heterozygous mice. Legs were shipped to USA, and used for the preparation of BMDCs within 24-48hr. All experimental procedures and protocols for mice are approved by the Animal Care and Facilities Committee at both Rutgers and University of Düsseldorf.

### Preparation of bone marrow-derived DCs (BMDCs) and spleen DCs

Preparation of mouse BMDCs was according to the method described in Inaba *et al*(90) with slight modification(24). Briefly, single cell suspension was prepared from bone marrow of femurs and tibias, and plated on 65mm dishes in Iscove’s modified Dulbecco’s Medium containing 10% fetal calf serum and 10ng/ml of GM-CSF for 7-8 days. Non-adherent cells were collected and DCs were purified by centrifugation over a 13.7% (w/v) metrizamide discontinuous gradient. More than 85% of cells collected at the interface of the gradient were positive for CD11c. Cells were matured by overnight culture by the addition of 100ng/ml of lipopolysaccharide (LPS, Sigma).

Spleen DCs were prepared essentially according to the protocol described in (91) with slight modifications. Briefly, a single cell suspension was prepared by treating spleens with collagenase. After overnight culture, dendritic cells were purified with two rounds of MACS magnetic separation using mouse CD11c microbeads (Miltenyi Biotec) according to the manufacturer’s instructions. More than 85% of cells are CD11c and MHC-II positive.

### Cytokine profiling

For *in vivo* cytokine profiling, WT and fascin1 KO mice were intravenously infected with *Listeria* monocytogenes (10403S, 1×10^4^ cfu/mouse). Two days later, sera were taken from euthanized mice and used for cytokine profiling with Mouse Cytokine Antibody Array C series 2000 (spanning 144 unique mouse cytokines/chemokines) according to the manufacturer’s instructions (RayBiotech). For analyses of cytokines in culture supernatants of BMDCs, WT and fascin1 KO BMDCs were cultured overnight after addition of LPS (100ng/ml). Culture supernatants were then used for profiling of inflammatory cytokines using a RayBio Quantibody mouse Th17 array1 according to the manufacturer’s instructions (RayBiotech). The Quantibody array allows us to quantitatively determine the concentrations of 18 cytokines/chemokines (IL-1β, IL-2, IL-4, IL-5, IL-6, IL-10, IL-12 p70, IL-17, IL17F, IL-21 IL-22, IL-23, IL-28, IFNγ, MIP-3α, TGFβ1, and TNFα). ELISA to determine IL-6 was performed using an ELISA kit from BD biosciences according to the manufacturer’s instructions.

### Proximity Ligation Assay (PLA) to determine IL-6Rα-gp130 association

PLA was performed according to the manufacturer’s instruction (O-link Bioscience, Uppsala, Sweden) with slight modification as described before(31). Briefly, BMDCs were fixed with 3.7% formaldehyde for 10min. Fixed BMDCs were incubated with anti-gp130 rabbit polyclonal and anti-IL-6Rα goat polyclonal antibodies. The rabbit and goat antibodies were labeled with anti-rabbit and antigoat PLA probes, respectively, each of which was conjugated with oligonucleotides. If they were within ~40nm, these nucleotides could be ligated to form a closed circle, which was amplified with DNA polymerase and detected with fluorescently labeled oligonucleotides.

### Modified Boyden chamber

Collagen-coated Transwell plates (3μm hole, Corning, Lowell, MA) were used for modified Boyden chamber chemotaxis assays(24). CCL19 (MIP3β), a chemokine for mature DCs, was added in bottom wells at the concentration of 0.6μg/ml. After addition of LPS (100ng/ml), WT and fascin1 KO BMDCs (2×10^5^ cells) were placed on top wells. To block IL-6 signaling, a neutralizing IL-6Rα antibody (AF1830, Goat polyclonal, R&D Systems) was added at a final concentration of 1μg/ml just before LPS addition. Normal goat IgG antibody was used as a control. After overnight incubation, cells migrated to the bottom wells were counted. In some experiments, the IL-6Rα antibody was added after overnight maturation of BMDCs with LPS.

Chemotaxis of IL-6Rα KO and WT (or heterozygous) BMDCs toward CCL19 was performed in the same way as above. To restore IL-6 signaling of IL-6Rα KO DCs, human IL-6 (15ng/ml, 206-IL, R&D Systems) and soluble IL-6Rα (sIL-6Rα, 25ng/ml, 227-SR, R&D Systems) were added 1hr after LPS addition.

### Live cell imaging and chemotaxis analyses

Chemotaxis of BMDCs toward CCL19 gradient in collagen gel was assayed according to Sixt and Lammermann(48). Briefly, BMDCs were mixed with rat tail collagen, Type I (unpepsinized native collagen, final concentration, 1.5mg/ml, ibidi, Fitchburg, WI) and the mixture was polymerized in a chamber. After addition of CCL19 (0.6μg/ml) at the top of the collagen gel, migration of BMDCs was observed by time-lapse microscopy at 2min intervals for 60-90min. Migration tracks were obtained by MTrackJ (imageJ)(92), and migration tracks, directness, migration speeds, and forward and perpendicular migration indexes were obtained by Chemotaxis and Migration Tool software (ibidi USA Wisconsin). Immovable DCs trapped in a collagen gel were not included for analyses.

### *In vivo* migration assays of BMDCs

BMDCs (1×10^6^ cells) were labeled with a cell Trace-CSFE dye (Thermo Fisher, final concentration, 2.5μM) according to the manufacturer’s instructions. CSFE-labeled BMDCs (total 100 μl) with 3μg of an antibody against IL-6Rα or a control normal goat antibody were injected subcutaneously into a dorsal aspect of a foot. One day later, lymph node cells were isolated from both popliteal and inguinal lymph nodes and CSFE-labeled DCs migrated into these lymph nodes were counted by flow cytometry. Total cell numbers of lymph nodes were also counted to determine the cellularity.

### Immunofluorescence & Western blots

Immunofluorescence of surface proteins including CCR7, IL-6Rα and gp130 was performed after fixation of 4% formaldehyde without permeabilization. Images were taken as Z-stacks (0.2μm spacing) with a DeltaVision Image Restoration Microscope system, deconvolved with the softWoRx software (Applied Precision Instruments). Projected images were generated with SoftWoRx or ImageJ (http://rsb.info.nih.gov/ij/). In some experiments, images were taken on a Nikon TE300 microscope with a 60x objective lens (NA 1.4) and presented without deconvolution. Exposure times for imaging, as well as the settings for deconvolution, were constant for all samples to be compared within any given experiment. For presentation, contrast and brightness of all images in a given figure were adjusted in the same way with Photoshop (Adobe).

To determine the levels of phopho-ERK1/2, as well as those of total ERK1/2, BMDCs before and after CCL19 treatment were homogenized in a lysis buffer containing 0.5% Triton-100X as described(93). After centrifugation, the supernatants were used for Western blotting with antibodies against phospho ERK1/2 and total ERK1/2 (Cell Signaling Technology).

### Flow cytometry analyses

For surface labeling of CCR7, gp130 and IL-6Rα, BMDCs were fixed with 3.7% formalin and incubated with a primary antibody to each protein. The primary antibodies were labeled with an FITC- or AF647-labeled second antibody. For other surface proteins including MHC-II, CD80, CD86, formalin-fixed BMDCs were labeled with a fluorescently-conjugated antibody to each protein. CD86-positive BMDCs were gated for flow cytometry analyses. Flow cytometry was performed with a BD FACSCalibur or Coulter Cytomics FC500 flow cytometer. We found that CCR7 staining of BMDCs is sometimes associated with cytoplasmic staining probably due to breakage of the dendritic membrane structure during staining procedures. To confirm membrane staining of CCR7, we also performed AMNIS imaging cytometry, which can specifically measure surface staining of CCR7.

### CCR7 internalization assay

To determine effects of IL-6 signaling on ligand-induced CCR7 internalization, BMDCs matured overnight with LPS were incubated with a neutralizing IL-6Rα antibody or control goat antibody for 10min. BMDCs were then treated with 0.6μg/ml of CCL19. BMDCs before and 5min after CCL19 activation were fixed with formalin, and labeled with anti-CCR7 and anti-CD86 antibodies. Flow cytometry was performed to determine changes in CCR7 surface expression upon CCL19 treatment.

Internalization of CCR7 was also examined with HEK293T cells stably expressing a mouse CCR7-GFP fusion construct (pCMV3-mCCR7-GFPSpark, Sino Biological Inc.). HEK293T cells were transfected with CCR7-GFP with Lipofectamin 2000 (Gibco). After selection with hygromycin, stable cell lines expressing a uniform level of CCR7-GFP were picked up with a small piece of a filter paper under a fluorescent microscope, and propagated in the presence hygromycin. To determine CCR7 internalization, CCR7-GFP-expressing HEC293T cells were incubated with or without a neutralizing antibody for 10min, then CCL19 (0.6μg/ml) was added for 5min. After fixing with 3.7% formaldehyde, cells were incubated with an anti-mouse CCR7 antibody premixed with a Cy3-labeled Fab fragment of the secondary antibody. Both GFP and Cy3 fluorescent images were taken by a Nikon TE300 or DeltaVision Image Restoration Microscope system. Cy3 fluorescent images represent surface expression of exogenously expressed mouse CCR7, which was quantitated by ImageJ.

### Statistical analysis

Statistical analyses were performed using a Student t test for two groups (http://www.physics.csbsju.edu/stats/t-test.html) or using ANOVA for more than 3 groups (Excel and StatPlus).

## Supporting information

video 1b with overlay

Video 2b with overlay

video 3b with overlay

## Acknowledgment

We wish to thank Dr. Ron P. Hart for his help to use his fluorescent glass slide reader, Dr. Michael Sixt for the help to set up chemotaxis in 3D collagen environment, Dr. Stefan Krautwald for CCL19-IgG fusion protein and Dr. D. A. Portnoy for *L. monocytogenes* strains.

## Footnotes

^#^This work was supported by R03AI099315 (to SY) and American Heart Association (to FM).

## Abbreviations

WT: wild type
KO: knockout
DCs: Dendritic cells
BM-DCs: Bone marrow-derived dendritic cells

## References

1. R. M. Steinman, H. Hemmi, Dendritic cells: translating innate to adaptive immunity. Curr Top Microbiol Immunol 311, 17–58 (2006).

2. I. Mellman, R. M. Steinman, Dendritic cells: specialized and regulated antigen processing machines. Cell 106, 255–258 (2001).

3. D. Alvarez, E. H. Vollmann, U. H. von Andrian, Mechanisms and consequences of dendritic cell migration. Immunity 29, 325–342 (2008).

4. I. J. De Vries et al., Effective migration of antigen-pulsed dendritic cells to lymph nodes in melanoma patients is determined by their maturation state. Cancer Res 63, 12–17 (2003).

5. M. C. Dieu et al., Selective recruitment of immature and mature dendritic cells by distinct chemokines expressed in different anatomic sites. J Exp Med 188, 373–386 (1998).

6. F. Sallusto et al., Rapid and coordinated switch in chemokine receptor expression during dendritic cell maturation. Eur J Immunol 28, 2760–2769 (1998).

7. S. Sozzani et al., Differential regulation of chemokine receptors during dendritic cell maturation: a model for their trafficking properties. J Immunol 161, 1083–1086 (1998).

8. S. Yanagihara, E. Komura, J. Nagafune, H. Watarai, Y. Yamaguchi, EBI1/CCR7 is a new member of dendritic cell chemokine receptor that is up-regulated upon maturation. J Immunol 161, 3096–3102 (1998).

9. L. Ohl et al., CCR7 governs skin dendritic cell migration under inflammatory and steady-state conditions. Immunity 21, 279–288 (2004).

10. R. Forster et al., CCR7 coordinates the primary immune response by establishing functional microenvironments in secondary lymphoid organs. Cell 99, 23–33 (1999).

11. R. Forster, A. C. Davalos-Misslitz, A. Rot, CCR7 and its ligands: balancing immunity and tolerance. Nat Rev Immunol 8, 362–371 (2008).

12. N. Sanchez-Sanchez et al., Chemokine receptor CCR7 induces intracellular signaling that inhibits apoptosis of mature dendritic cells. Blood 104, 619–625 (2004).

13. G. Bardi, V. Niggli, P. Loetscher, Rho kinase is required for CCR7-mediated polarization and chemotaxis of T lymphocytes. FEBS Lett 542, 79–83 (2003).

14. M. A. Allaire, N. Dumais, Involvement of the MAPK and RhoA/ROCK pathways in PGE2-mediated CCR7-dependent monocyte migration. Immunology letters 146, 70–73 (2012).

15. Y. Yanagawa, K. Onoe, CCR7 ligands induce rapid endocytosis in mature dendritic cells with concomitant up-regulation of Cdc42 and Rac activities. Blood 101, 4923–4929 (2003).

16. J. V. Stein et al., CCR7-mediated physiological lymphocyte homing involves activation of a tyrosine kinase pathway. Blood 101, 38–44 (2003).

17. L. Riol-Blanco et al., The chemokine receptor CCR7 activates in dendritic cells two signaling modules that independently regulate chemotaxis and migratory speed. J Immunol 174, 4070–4080 (2005).

18. J. C. Adams, Roles of fascin in cell adhesion and motility. Curr Opin Cell Biol 16, 590–596 (2004).

19. A. Jayo, M. Parsons, Fascin: a key regulator of cytoskeletal dynamics. Int J Biochem Cell Biol 42, 1614–1617 (2010).

20. S. Yamashiro, Functions of fascin in dendritic cells. Critical reviews in immunology 32, 11–21 (2012).

21. L. Sonderbye, R. Magerstadt, R. N. Blatman, F. I. Preffer, E. Langhoff, Selective expression of human fascin (p55) by dendritic leukocytes. Adv Exp Med Biol 417, 41–46 (1997).

22. R. Ross, X. L. Ross, J. Schwing, T. Langin, A. B. Reske-Kunz, The actin-bundling protein fascin is involved in the formation of dendritic processes in maturing epidermal Langerhans cells. J Immunol 160, 3776–3782 (1998).

23. G. Mosialos et al., Circulating human dendritic cells differentially express high levels of a 55-kd actin-bundling protein. Am J Pathol 148, 593–600 (1996).

24. Y. Yamakita et al., Fascin1 promotes cell migration of mature dendritic cells. J Immunol 186, 2850–2859 (2011).

25. S. Yamashiro, Y. Yamakita, S. Ono, F. Matsumura, Fascin, an actin-bundling protein, induces membrane protrusions and increases cell motility of epithelial cells. Mol. Biol. Cell 9, 993–1006 (1998).

26. L. Chen, S. Yang, J. Jakoncic, J. J. Zhang, X. Y. Huang, Migrastatin analogues target fascin to block tumour metastasis. Nature 464, 1062–1066 (2010).

27. J. Yao et al., Signal transducer and activator of transcription 3 signaling upregulates fascin via nuclear factor-kappaB in gastric cancer: Implications in cell invasion and migration. Oncology letters 7, 902–908 (2014).

28. D. Vignjevic et al., Fascin, a novel target of beta-catenin-TCF signaling, is expressed at the invasive front of human colon cancer. Cancer Res 67, 6844–6853 (2007).

29. Y. Hashimoto, M. Parsons, J. C. Adams, Dual actin-bundling and protein kinase C-binding activities of fascin regulate carcinoma cell migration downstream of Rac and contribute to metastasis. Mol. Biol. Cell 18, 4591–4602 (2007).

30. F. K. Huang et al., Targeted inhibition of fascin function blocks tumour invasion and metastatic colonization. Nat Commun 6, 7465 (2015).

31. F. Matsumura, Y. Yamakita, V. Starovoytov, S. Yamashiro, Fascin confers resistance to Listeria infection in dendritic cells. J Immunol 191, 6156–6164 (2013).

32. F. Li et al., miR-98 suppresses melanoma metastasis through a negative feedback loop with its target gene IL-6. Experimental & molecular medicine 46, e116 (2014).

33. S. M. Hurst et al., Il-6 and its soluble receptor orchestrate a temporal switch in the pattern of leukocyte recruitment seen during acute inflammation. Immunity 14, 705–714 (2001).

34. H. Jayatilaka et al., Synergistic IL-6 and IL-8 paracrine signalling pathway infers a strategy to inhibit tumour cell migration. Nat Commun 8, 15584 (2017).

35. W. C. Wang, C. Y. Kuo, B. S. Tzang, H. M. Chen, S. H. Kao, IL-6 augmented motility of airway epithelial cell BEAS-2B via Akt/GSK-3beta signaling pathway. J Cell Biochem 113, 3567–3575 (2012).

36. G. X. Jiang et al., IL-6/STAT3/TFF3 signaling regulates human biliary epithelial cell migration and wound healing in vitro. Mol Biol Rep 37, 3813–3818 (2010).

37. J. K. Kim, S. M. Lee, K. Suk, W. H. Lee, A novel pathway responsible for lipopolysaccharide-induced translational regulation of TNF-alpha and IL-6 expression involves protein kinase C and fascin. J Immunol 187, 6327–6334 (2011).

38. M. Hibi et al., Molecular cloning and expression of an IL-6 signal transducer, gp130. Cell 63, 1149–1157 (1990).

39. A. E. Morelli et al., Cytokine production by mouse myeloid dendritic cells in relation to differentiation and terminal maturation induced by lipopolysaccharide or CD40 ligation. Blood 98, 1512–1523 (2001).

40. D. Nagorsen, F. M. Marincola, M. C. Panelli, Cytokine and chemokine expression profiles of maturing dendritic cells using multiprotein platform arrays. Cytokine 25, 31–35 (2004).

41. J. Helft et al., GM-CSF Mouse Bone Marrow Cultures Comprise a Heterogeneous Population of CD11c(+)MHCII(+) Macrophages and Dendritic Cells. Immunity 42, 1197–1211 (2015).

42. D. Jin, J. Sprent, GM-CSF Culture Revisited: Preparation of Bulk Populations of Highly Pure Dendritic Cells from Mouse Bone Marrow. J Immunol 201, 3129–3139 (2018).

43. M. B. Lutz, K. Inaba, G. Schuler, N. Romani, Still Alive and Kicking: In-Vitro-Generated GM-CSF Dendritic Cells!Immunity 44, 1–2 (2016).

44. M. Guilliams, B. Malissen, A Matter of Perspective: Moving from a Pre-omic to a Systems-Biology Vantage of Monocyte-Derived Cell Function and Nomenclature. Immunity 44, 5–6 (2016).

45. M. Guilliams, B. Malissen, A Death Notice for In-Vitro-Generated GM-CSF Dendritic Cells? Immunity 42, 988–990 (2015).

46. M. B. Lutz, H. Strobl, G. Schuler, N. Romani, GM-CSF Monocyte-Derived Cells and Langerhans Cells As Part of the Dendritic Cell Family. Frontiers in immunology 8, 1388 (2017).

47. Y. Xu, Y. Zhan, A. M. Lew, S. H. Naik M. H. Kershaw, Differential development of murine dendritic cells by GM-CSF versus Flt3 ligand has implications for inflammation and trafficking. J Immunol 179, 7577–7584 (2007).

48. M. Sixt, T. Lammermann, In vitro analysis of chemotactic leukocyte migration in 3D environments. Methods Mol Biol 769, 149–165 (2011).

49. E. F. Foxman, E. J. Kunkel, E. C. Butcher, Integrating conflicting chemotactic signals. The role of memory in leukocyte navigation. J Cell Biol 147, 577–588 (1999).

50. P. Zengel et al., mu-Slide Chemotaxis: a new chamber for long-term chemotaxis studies. BMC Cell Biol 12, 21 (2011).

51. T. Lammermann et al., Rapid leukocyte migration by integrin-independent flowing and squeezing. Nature 453, 51–55 (2008).

52. C. A. Hunter, S. A. Jones, IL-6 as a keystone cytokine in health and disease. Nat Immunol 16, 448–457 (2015).

53. S. Rose-John, M. F. Neurath, IL-6 trans-signaling: the heat is on. Immunity 20, 2–4 (2004).

54. S. A. Jones, S. Rose-John, The role of soluble receptors in cytokine biology: the agonistic properties of the sIL-6R/IL-6 complex. Biochim Biophys Acta 1592, 251–263 (2002).

55. F. A. Montero-Julian, The soluble IL-6 receptors: serum levels and biological function. Cell Mol Biol (Noisy-le-grand) 47, 583–597 (2001).

56. S. A. Jones, S. Horiuchi, N. Topley, N. Yamamoto, G. M. Fuller, The soluble interleukin 6 receptor: mechanisms of production and implications in disease. FASEB J 15, 43–58 (2001).

57. G. H. Fan, W. Yang, X. J. Wang, Q. Qian, A. Richmond, Identification of a motif in the carboxyl terminus of CXCR2 that is involved in adaptin 2 binding and receptor internalization. Biochemistry 40, 791–800 (2001).

58. K. Schaeuble et al., Ubiquitylation of the chemokine receptor CCR7 enables efficient receptor recycling and cell migration. J Cell Sci 125, 4463–4474 (2012).

59. Y. Daaka et al., Essential role for G protein-coupled receptor endocytosis in the activation of mitogen-activated protein kinase. J Biol Chem 273, 685–688 (1998).

60. S. Krautwald et al., Ectopic expression of CCL19 impairs alloimmune response in mice. Immunology 112, 301–309 (2004).

61. G. F. Debes et al., Chemokine receptor CCR7 required for T lymphocyte exit from peripheral tissues. Nat Immunol 6, 889–894 (2005).

62. W. Zhao, M. Liu, K. L. Kirkwood, p38alpha stabilizes interleukin-6 mRNA via multiple AU-rich elements. J Biol Chem 283, 1778–1785 (2008).

63. S. Paschoud et al., Destabilization of interleukin-6 mRNA requires a putative RNA stem-loop structure, an AU-rich element, and the RNA-binding protein AUF1. Mol Cell Biol 26, 8228–8241 (2006).

64. K. Masuda et al., Arid5a controls IL-6 mRNA stability, which contributes to elevation of IL-6 level in vivo. Proc. Natl. Acad. Sci. USA 110, 9409–9414 (2013).

65. C. Y. Brown, C. A. Lagnado, M. A. Vadas, G. J. Goodall, Differential regulation of the stability of cytokine mRNAs in lipopolysaccharide-activated blood monocytes in response to interleukin-10. J Biol Chem 271, 20108–20112 (1996).

66. Y. Seko, S. Cole, W. Kasprzak, B. A. Shapiro, J. A. Ragheb, The role of cytokine mRNA stability in the pathogenesis of autoimmune disease. Autoimmunity reviews 5, 299–305 (2006).

67. C. Otero, M. Groettrup, D. F. Legler, Opposite fate of endocytosed CCR7 and its ligands: recycling versus degradation. J Immunol 177, 2314–2323 (2006).

68. H. Arai, F. S. Monteclaro, C. L. Tsou, C. Franci, I. F. Charo, Dissociation of chemotaxis from agonist-induced receptor internalization in a lymphocyte cell line transfected with CCR2B. Evidence that directed migration does not require rapid modulation of signaling at the receptor level. J Biol Chem 272, 25037–25042 (1997).

69. M. A. Byers et al., Arrestin 3 mediates endocytosis of CCR7 following ligation of CCL19 but not CCL21. J Immunol 181, 4723–4732 (2008).

70. T. A. Kohout et al., Differential desensitization, receptor phosphorylation, beta-arrestin recruitment, and ERK1/2 activation by the two endogenous ligands for the CC chemokine receptor 7. J Biol Chem 279, 23214–23222 (2004)

71. D. A. Zidar, J. D. Violin, E. J. Whalen, R. J. Lefkowitz, Selective engagement of G protein coupled receptor kinases (GRKs) encodes distinct functions of biased ligands. Proc. Natl. Acad. Sci. USA 106, 9649–9654 (2009).

72. S. J. Park et al., IL-6 regulates in vivo dendritic cell differentiation through STAT3 activation. J Immunol 173, 3844–3854 (2004).

73. V. N. Goncharuk, J. S. Ross, J. A. Carlson, Actin-binding protein fascin expression in skin neoplasia. J Cutan Pathol 29, 430–438 (2002).

74. A. Grothey et al., C-erbB-2/HER-2 upregulates fascin, an actin-bundling protein associated with cell motility, in human breast cancer cell lines. Oncogene 19, 4864–4875 (2000).

75. W. Hu et al., Increased expression of fascin, motility associated protein, in cell cultures derived from ovarian cancer and in borderline and carcinomatous ovarian tumors. Clin Exp Metastasis 18, 83–88 (2000).

76. L. M. Machesky, A. Li, Fascin: Invasive filopodia promoting metastasis. Commun Integr Biol 3, 263–270 (2010).

77. A. Peraud et al., Expression of fascin, an actin-bundling protein, in astrocytomas of varying grades. Brain Tumor Pathol 20, 53–58 (2003).

78. S. Sun et al., Expression of fascin in breast cancer correlates with tumor grade, increased cell proliferation and p53 tumor suppressor gene overexpression. Laboratory Investigation 76, 26A (1997).

79. A. Li et al., The actin-bundling protein fascin stabilizes actin in invadopodia and potentiates protrusive invasion. Curr Biol 20, 339–345 (2010).

80. J. Bollrath et al., gp130-mediated Stat3 activation in enterocytes regulates cell survival and cell-cycle progression during colitis-associated tumorigenesis. Cancer Cell 15, 91–102 (2009).

81. A. Ferraresi et al., Resveratrol inhibits IL-6-induced ovarian cancer cell migration through epigenetic up-regulation of autophagy. Mol Carcinog 10.1002/mc.22582 (2016).

82. L. R. Luckett, R. M. Gallucci, Interleukin-6 (IL-6) modulates migration and matrix metalloproteinase function in dermal fibroblasts from IL-6KO mice. Br J Dermatol 156, 1163–1171 (2007).

83. W. Sun et al., Interleukin-6 promotes the migration and invasion of nasopharyngeal carcinoma cell lines and upregulates the expression of MMP-2 and MMP-9. Int J Oncol 44, 1551–1560 (2014).

84. I. Tamm et al., Interleukin 6 decreases cell-cell association and increases motility of ductal breast carcinoma cells. J Exp Med 170, 1649–1669 (1989).

85. M. Walter, S. Liang, S. Ghosh, P. J. Hornsby, R. Li, Interleukin 6 secreted from adipose stromal cells promotes migration and invasion of breast cancer cells. Oncogene 28, 2745–2755 (2009).

86. S. Grivennikov et al., IL-6 and Stat3 are required for survival of intestinal epithelial cells and development of colitis-associated cancer. Cancer Cell 15, 103–113 (2009).

87. Y. K. Mburu et al., Chemokine receptor 7 (CCR7) gene expression is regulated by NF-kappaB and activator protein 1 (AP1) in metastatic squamous cell carcinoma of head and neck (SCCHN). J Biol Chem 287, 3581–3590 (2012).

88. Y. Yamakita, F. Matsumura, S. Yamashiro, Fascin1 is dispensable for mouse development but is favorable for neonatal survival. Cell Motil Cytoskeleton 66, 524–534 (2009).

89. M. M. McFarland-Mancini et al., Differences in wound healing in mice with deficiency of IL-6 versus IL-6 receptor. J Immunol 184, 7219–7228 (2010).

90. K. Inaba et al., Generation of large numbers of dendritic cells from mouse bone marrow cultures supplemented with granulocyte/macrophage colonystimulating factor. J Exp Med 176, 1693–1702 (1992).

91. A. J. Stagg, F. Burke, S. Hill, S. C. Knight, “isolation of mouse spleen dendritic cells” in Dendritic cell protocols, S. P. Robinson, A. J. Stagg, Eds. (humana Press, New Jersey, 2001), vol. 64, pp. 9–22.

92. E. Meijering, O. Dzyubachyk, I. Smal, Methods for cell and particle tracking. Methods Enzymol 504, 183–200 (2012).

93. S. Yamashiro et al., Myosin phosphatase-targeting subunit 1 regulates mitosis by antagonizing polo-like kinase 1. Dev Cell 14, 787–797 (2008).

