## Supplemental Figures and video for "Investigation of fascin1, a marker of mature dendritic cells, reveals a New role for IL-6 signaling in chemotaxis"

**Supplemental Information**:

### Supplemental Fig. 1

**A**

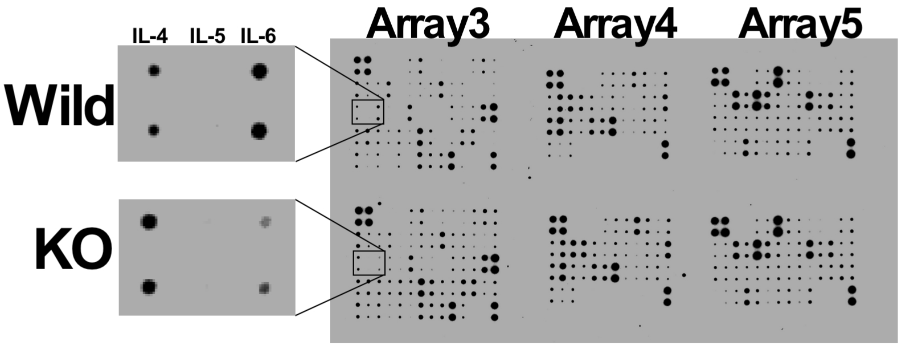


**B**

#### RayBio® Mouse Cytokine Antibody Array 3 (62)

| | A | B | C | D | E | F | G | H | I | J | K | L | M | N |
|---|---|---|---|---|---|---|---|---|---|---|---|---|---|---|
| 1 | POS | POS | NEG | NEG | Blank | Axl | BLC | CD30 L | CD30 T | CD40 | CRG-2 | CTACK | CXCL16 | Eotaxin |
| 2 | POS | POS | NEG | NEG | Blank | Axl | BLC | CD30 L | CD30 T | CD40 | CRG-2 | CTACK | CXCL16 | Eotaxin |
| 3 | Eotaxin-2 | Fas Ligand | Fractalkine | GCSF | GM-CSF | IFN γ | IGFBP-3 | IGFBP-5 | IGFBP-6 | IL-1α | IL-1 beta | IL-2 | IL-3 | IL-3 Rb |
| 4 | Eotaxin-2 | Fas Ligand | Fractalkine | GCSF | GM-CSF | IFN γ | IGFBP-3 | IGFBP-5 | IGFBP-6 | IL-1α | IL-1 beta | IL-2 | IL-3 | IL-3 Rb |
| 5 | IL-4 | IL-5 | IL-6 | IL-9 | IL-10 | IL-12 p40/p70 | IL-12 p70 | IL-13 | IL-17 | KC | Leptin R | Leptin | LIX | L-Selectin |
| 6 | IL-4 | IL-5 | IL-6 | IL-9 | IL-10 | IL-12 p40/p70 | IL-12 p70 | IL-13 | IL-17 | KC | Leptin R | Leptin | LIX | L-Selectin |
| 7 | Lymphotactin | MCP1 | MCP-5 | M-CSF | MIG | MIP-1α | MIP-1γ | MIP-2 | MIP-3 β | MIP-3α | PF-4 | P-Selectin | RANTES | SCF |
| 8 | Lymphotactin | MCP1 | MCP-5 | M-CSF | MIG | MIP-1α | MIP-1γ | MIP-2 | MIP-3 β | MIP-3α | PF-4 | P-Selectin | RANTES | SCF |
| 9 | SDF-1α | TARC | TCA-3 | TECK | TIMP-1 | TNFα | sTNF RI | sTNF RII | TPO | VCAM-1 | VEGF | Blank | Blank | POS |
| # | SDF-1α | TARC | TCA-3 | TECK | TIMP-1 | TNFα | sTNF RI | sTNF RII | TPO | VCAM-1 | VEGF | Blank | Blank | POS |

#### RayBio® Mouse Cytokine Antibody Array 4 (34)

| | A | B | C | D | E | F | G | H | I | J | K | L |
|---|---|---|---|---|---|---|---|---|---|---|---|---|
| 1 | POS | POS | NEG | NEG | BLANK | bFGF | DPPIV/CD26 | Dtk | E-Selectin | Fcγ RIIB | lt-3 Ligan | GITR |
| 2 | POS | POS | NEG | NEG | BLANK | bFGF | DPPIV/CD26 | Dtk | E-Selectin | Fcγ RIIB | lt-3 Ligan | GITR |
| 3 | HGF R | ICAM-1 | IGFBP-2 | IGF-I | IGF-II | IL-15 | IL-17B R | IL-7 | I-TAC | Lungkine | MDC | MMP-2 |
| 4 | HGF R | ICAM-1 | IGFBP-2 | IGF-I | IGF-II | IL-15 | IL-17B R | IL-7 | I-TAC | Lungkine | MDC | MMP-2 |
| 5 | MMP-3 | Osteopontin | Osteoporotegerin | Pro-MMP-9 | Resistin | Shh-N | Thymus CK-1 | TIMP-2 | TRANCE | TROY | TSLP | VEGF R1 |
| 6 | MMP-3 | Osteopontin | Osteoporotegerin | Pro-MMP-9 | Resistin | Shh-N | Thymus CK-1 | TIMP-2 | TRANCE | TROY | TSLP | VEGF R1 |
| 7 | VEGF R2 | VEGF R3 | VEGF-D | BLANK | BLANK | BLANK | BLANK | BLANK | BLANK | BLANK | BLANK | POS |
| 8 | VEGF R2 | VEGF R3 | VEGF-D | BLANK | BLANK | BLANK | BLANK | BLANK | BLANK | BLANK | BLANK | POS |

#### RayBio® Mouse Cytokine Antibody Array 5 (48)

| | A | B | C | D | E | F | G | H | I | J | K | L | M | N |
|---|---|---|---|---|---|---|---|---|---|---|---|---|---|---|
| 1 | POS | POS | NEG | NEG | 4-1BB | 6Ckine | ACE /CD143 | ALK-1 | Amphiregulin | Cardiotrophin-1 | CD27 | CD27 Ligand | CD36 | CD40 Ligand |
| 2 | POS | POS | NEG | NEG | 4-1BB | 6Ckine | ACE /CD143 | ALK-1 | Amphiregulin | Cardiotrophin-1 | CD27 | CD27 Ligand | CD36 | CD40 Ligand |
| 3 | Chordin | CTLA-4 | Decorin | DKK-1 | E-Cadherin | EGF | Endoglin | Epigen | Epiregulin | Galectin-1 | Growth arrest specific 1 | Growth arrest specific 6 | GITR Ligand | Granzyme B |
| 4 | Chordin | CTLA-4 | Decorin | DKK-1 | E-Cadherin | EGF | Endoglin | Epigen | Epiregulin | Galectin-1 | Growth arrest specific 1 | Growth arrest specific 6 | GITR Ligand | Granzyme B |
| 5 | HAI-1 | HGF | IL-1 R4/ST2L | IL-11 | IL-17B | IL-17E | IL-17F | IL-1ra/IL-1F3 | IL-2 R alpha | IL-20 | IL-21 | IL-28 | IL-6 R | JAM-A |
| 6 | HAI-1 | HGF | IL-1 R4/ST2L | IL-11 | IL-17B | IL-17E | IL-17F | IL-1ra/IL-1F3 | IL-2 R alpha | IL-20 | IL-21 | IL-28 | IL-6 R | JAM-A |
| 7 | MAdCAM-1 | MFG-E8 | Neprilysin | Pentraxin 3 | Prolactin | RAGE | TACI | TREM-1 | TWEAK | TWEAK R | NEG | NEG | NEG | POS |
| 8 | MAdCAM-1 | MFG-E8 | Neprilysin | Pentraxin 3 | Prolactin | RAGE | TACI | TREM-1 | TWEAK | TWEAK R | NEG | NEG | NEG | POS |

Abbreviations: Pos: positive control; Neg: negative control. All others use standard abbreviations.

Note: IL-12 reacts both IL-12p40 and IL-12p70. IL-12p70 only recognizes IL-12p70.

**Supplemental Figure 1.** Analysis of cytokine profiles in sera of wild type and fascin1 KO mice. Mice were infected with *Listeria monocytogenes*. Sera taken 48hr after infection were analyzed by a RayBio Mouse cytokine antibody Arrays 3-5. (**A**) Dot blot showing specific reduction of IL-6 in fascin1 KO mouse sera. (**B**) Panels of cytokine/chemokines in arrays 1-3 are shown.

Supplemental Figure 2

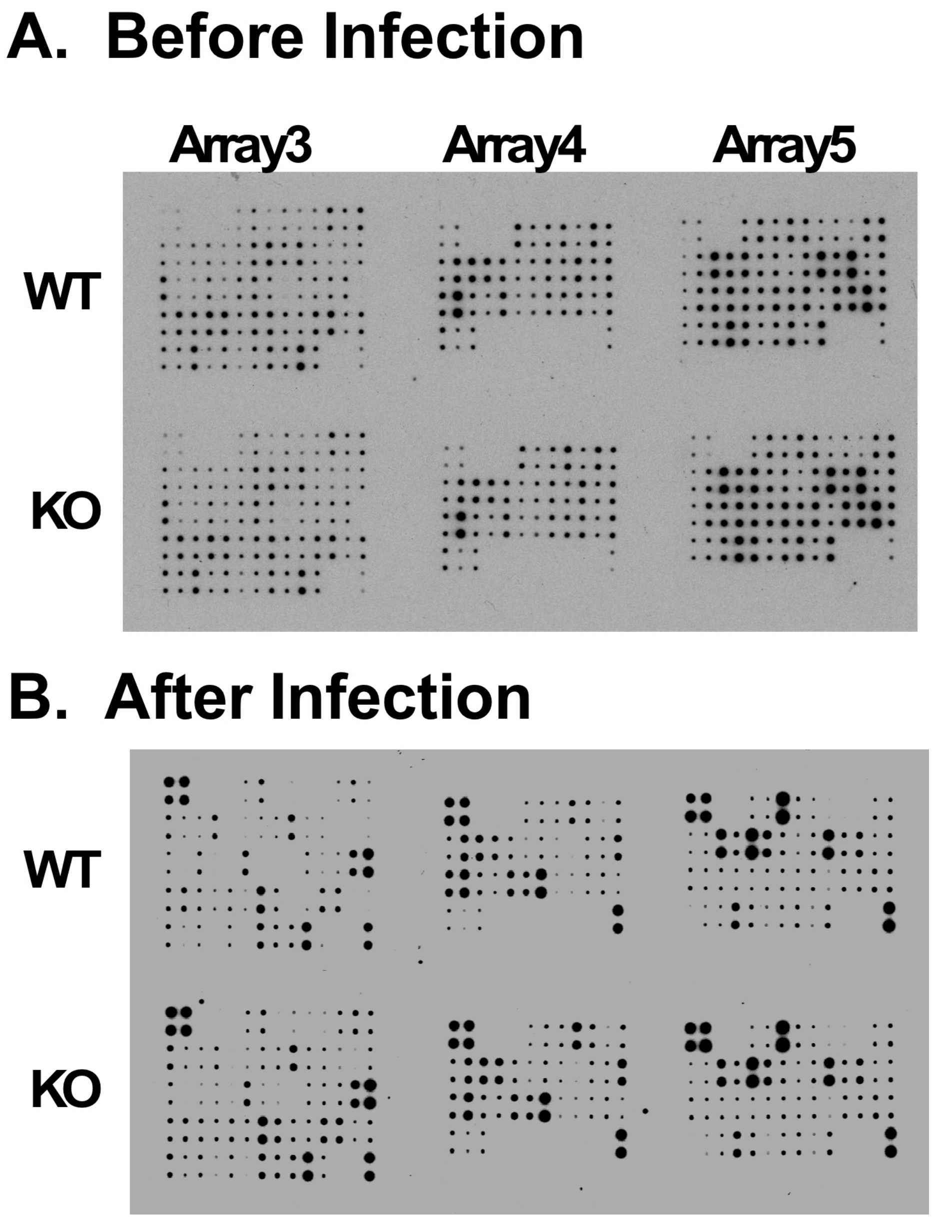


**Supplemental Figure 2**. Cytokine profiles of sera before and after infection with *Listeria monocytogenes*. The profile was determined by a RayBio Mouse cytokine antibody Arrays 3-5.

#### Supplemental Fig. 3

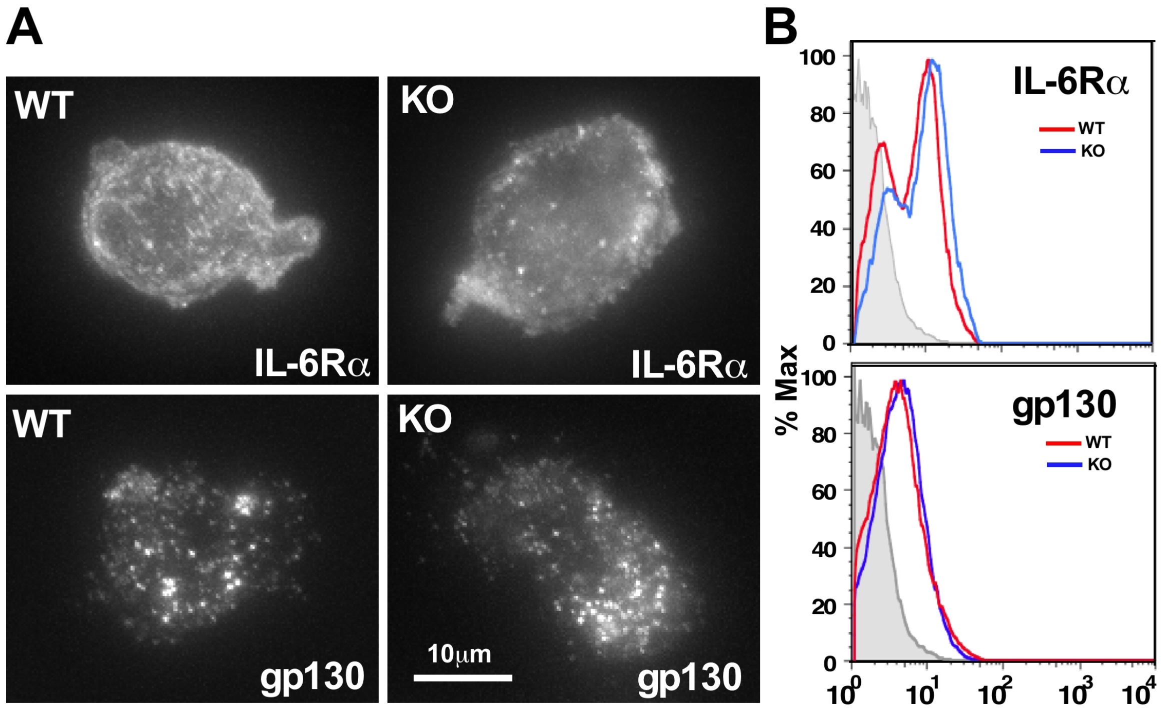


**Supplemental Figure 3**. Localization and surface expression of IL-6Rα and gp130 were similar between WT and fascin1 KO BMDCs. A, Immunofluorescent localization of IL-6Rα and gp130 in WT and fascin1 KO BMDCs. B, Flow cytometry analyses of surface expression of IL-6Rα and gp130 in WT and fascin1 KO BMDCs. The flow cytometry analysis of surface IL-6Rα (D) indicates two populations of BMDCs with high and low IL-6Rα, which may suggest two different subtypes of BMDCs.

#### Supplemental Figure 4

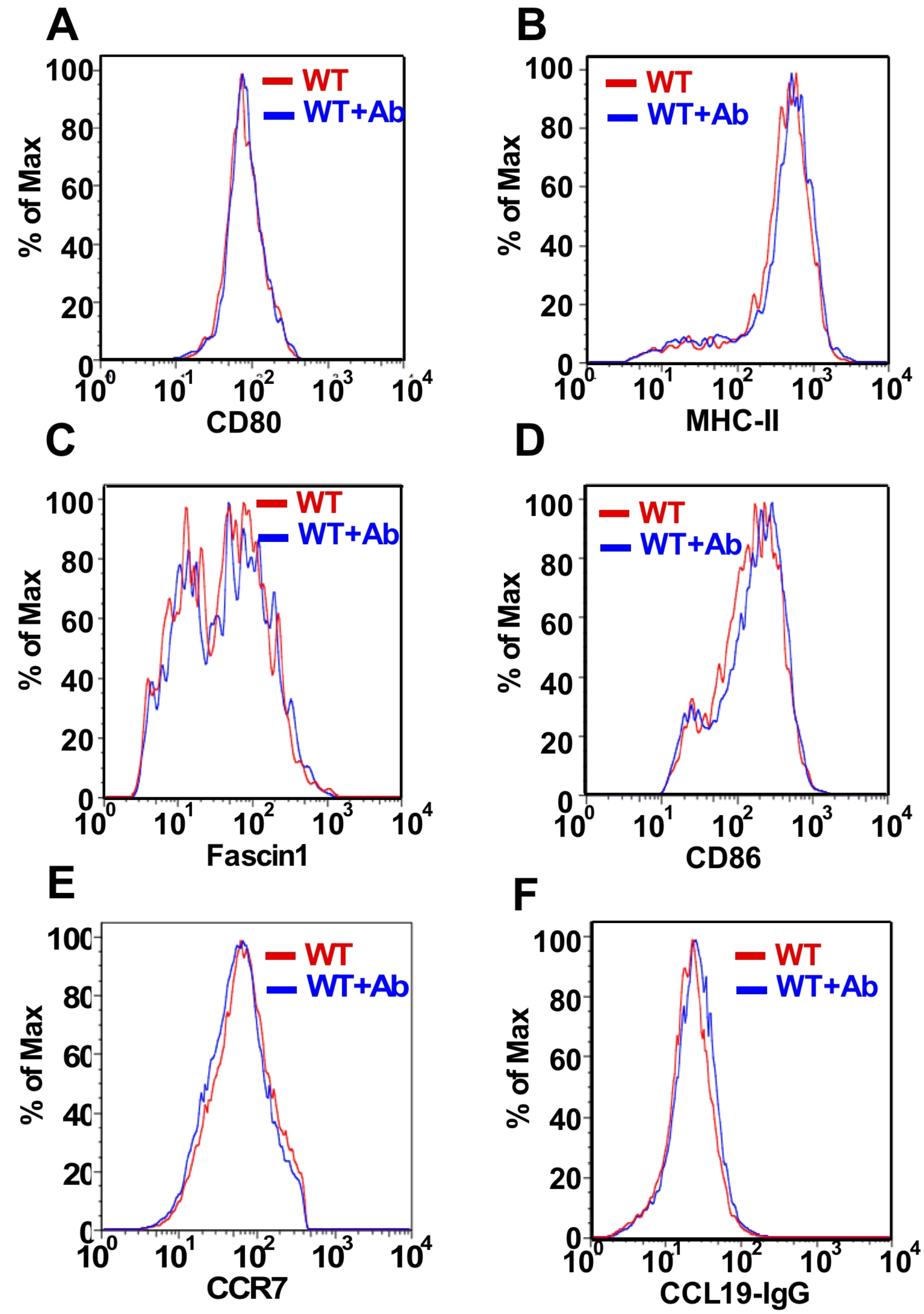


**Supplemental Figure 4.** Effects of the blockage of IL-6 signaling on the expression of DC maturation markers and on the association between CCR7 and CCL19. BMDCs were matured overnight by LPS in the presence or absence of a neutralizing antibody to IL-6Rα. Surface expression of maturation markers was determined by flow cytometry. A, CD80; B, MHC-II; C, fascin1; D, CD86; E, CCR7. The expression of these maturation markers is similar with or without neutralizing antibody against IL-6Rα. F, The association between CCR7 and CCL19 was examined using a CCL19-human IgG Fc fusion protein, followed by FITC-labeled anti-human IgG. No significant difference was observed.

**Supplemental Figure 5**

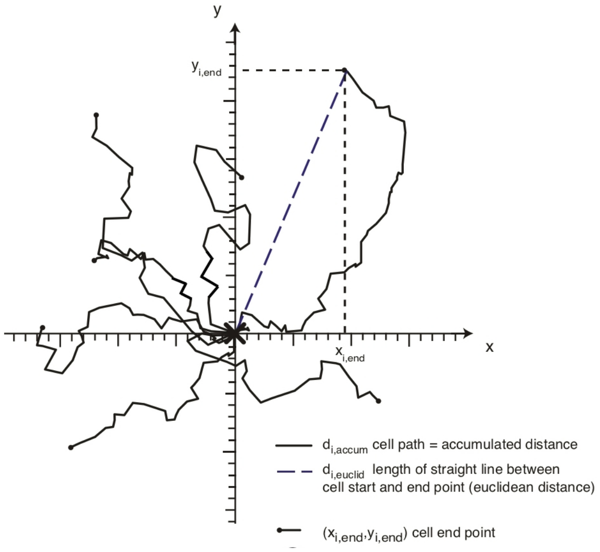


$$FMI_{\parallel} = \frac{y_{i,end}}{d_{i,accum}}$$

$$FMI_{\perp} = \frac{x_{i,end}}{d_{i,accum}}$$

$$Directness = \frac{d_{i,euclid}}{d_{i,accum}}$$

**Supplemental Figure 5.** Definition of parallel ($FMI_{\parallel}$) and perpendicular ($FMI_{\parallel}$) forward migration indexes and Directness of each migrating cell. $d_{i,\ accum}$, the entire migration path length of $i^{th}$ cell; $d_{i,\ euclid}$, euclidean length of straight line from the origin to the end point of $i^{th}$ cell. The figure is adopted from ref (1).

**Supplemental videos:**

Live cell imaging of BMDCs in a collagen gel under CCL19 gradient. All videos were taken with an upward CCL19 gradient. One second represents 20min. Scale bar, 50μm.

**Video 1a** WT BMDCs in a collagen gel under CCL19 gradient.

**Video 1b** The same video of 1a with overlay of a dot and line representing the cell body and migrating path of each migrating cell, respectively.

**Video 2a** Live cell imaging of fascin1 KO BMDCs in a collagen gel under CCL19 gradient.

**Video 2b** The same video of 2a with overlay of a dot and line representing the cell body and migrating path of each migrating cell, respectively.

**Video 3a** Live cell imaging of wild type BMDCs in a collagen gel under CCL19 gradient when IL-6 signaling is blocked with a function-inhibiting antibody against IL-6Rα.

**Video 3b** The same video of 3a with overlay of a dot and line representing the cell body and migrating path of each migrating cell, respectively.

**References:**

1. P. Zengel *et al.*, mu-Slide Chemotaxis: a new chamber for long-term chemotaxis studies. *BMC Cell Biol* **12**, 21 (2011).
